# Cytokinetic abscission is part of the mid-blastula transition switch in early zebrafish embryogenesis

**DOI:** 10.1101/2020.07.26.221515

**Authors:** Shai Adar-Levor, Dikla Nachmias, Shani T. Gal-Oz, Yarden M. Jahn, Nadine Peyrieras, Assaf Zaritsky, Ramon Y. Birnbaum, Natalie Elia

## Abstract

Animal cytokinesis ends with the formation of a thin intercellular membrane bridge connecting the two newly formed sibling cells that is ultimately resolved by abscission. While mitosis is completed within 15 minutes, the intercellular bridge can persist for hours, maintaining a physical connection between sibling cells and allowing exchange of cytosolic components. Although cell-cell communication is fundamental for development, the potential role of intercellular bridges during embryogenesis have not been fully elucidated. Here, we found that in early zebrafish (*Danio rerio*) embryogenesis, abscission is delayed and cells do not resolve their intercellular bridges until midblastula transition (MBT), giving rise to the formation of small inter-connected cell clusters. Interestingly, abscission commences during the MBT switch, which is manifested by cell cycle elongation, loss of synchronized divisions and genome activation. Moreover, depletion of Chmp4bb which is an essential ESCRT-III component for scission, delayed abscission beyond the MBT switch. Hallmark features of MBT, including transcription onset and cell shape changes, were similar in sibling cells connected by intercellular bridges, proposing a role for intercellular bridges in maintaining cell-cell communication in the embryo. Taken together, our data suggest that abscission is part of the cellular changes that occur during MBT and that cells coordinate their behavior during this critical embryonic phase through persisted intercellular bridges.

**Significance Statement:** In this work we show that the last step of cytokinesis, termed abscission, is inhibited in early zebrafish embryos. As a result, sibling cells remain connected to one another for several cycles and mutually time their developmental progress including transcription onset. Abscission commences at the 10^th^ cell cycle, when embryos enter the midblastula transition (MBT) switch in which embryonic cells become individualized and exhibit the characteristics of mature cells. Our data suggest that abscission is part of the MBT switch and that embryonic sibling cells mutually time their developmental progress by maintaining physical connections between them in the early embryo.

## INTRODUCTION

Cytokinesis is the part of cell division during which the cytoplasm of a single eukaryotic cell divides into two sibling cells. Animal cytokinesis begins with furrow ingression and is followed by the formation of a thin (~1 μm) and long (~10 μm) intercellular bridge that is packed with dense microtubule stalks and is surrounded by membrane (1–3). The intercellular bridge is ultimately severed in a process termed abscission that leads to the physical separation of the two daughter cells (1, 4). Abscission is a highly coordinated process that is executed by the ESCRT membrane fission machine and the microtubule-severing enzyme, Spastin, and is temporally regulated by cytokinetic proteins such as Aurora B (4–7).

The intercellular bridge persists for different durations, ranging from minutes to hours, depending on the cell type cellular context (2, 8–11). During this period, the cells are connected to one another and are able to exchange cytosolic components until abscission commences and the two cells become physically separated (10–13). Although chronologically abscission is the last step of cell division, it was previously shown that cells can continue to advance in cell cycle without resolving their intercellular bridges and that in mammalian cells, abscission commences at the G1 phase of the following cell cycle (7, 8). The ability to maintain cell connectivity while progressing through cell cycle, which can be mediated by regulating abscission timing, may have implications during early stages of development. Yet, although mitosis in general and cytokinesis in particular have been extensively studied during development, the role of abscission itself in a developmental setup has not been fully elucidated (3, 14–17).

The molecular components that drive and regulate abscission in *C.elegans* embryogenesis and in *Drosophila* oogenesis were shown to be similar to those documented in mammalian cells. However, the spatiotemporal characteristics of abscission and its regulation appeared to be different between these model systems and, in some cases, was associated with specific developmental phenotypes (7, 9, 10, 18). For example, in *Drosophila* germ cells, abscission delay led to the formation of stem-cysts, where all cells share the same cytoplasm but each cell remained individualized (10). Furthermore, abscission timing is differentially regulated in

*Drosophila* germ cells and interfering with it caused developmental abnormalities, such as encapsulation defects (10, 18). Additionally, in zebrafish (*Danio rerio*) oocytes, *C. elegans* embryos and MDCK (Madin-Darby canine kidney) cysts, the position of the intercellular bridge and the timing of abscission were directly related to cell polarization (7, 19–21). Together, these findings suggest that abscission may play a role in regulating early developmental processes.

Zebrafish embryogenesis begins with 10 cycles of rapid (15 minutes), synchronized divisions, composed of mitotic (M) and synthesis (S) phases while missing the gap phases (G1 and G2) (22, 23). At the end of the fourth cell cycle (16-cell stage), complete plasma membrane furrowing occurs for the first time and intercellular bridges can be formed (23). During the fast cell cycles, transcription and protein synthesis is inhibited. As a consequence, cell volume decreases in each cycle and cellular behavior is mediated by maternally deposited mRNAs and proteins, which are diluted from one cycle to the other (22, 24). At the 10^th^ division cycle, midblastula transition (MBT) occurs. The MBT is characterized by cell cycle lengthening (including gap phases), loss of cell synchrony, apoptosis, appearance of cell motility and onset of zygotic transcription (called zygotic genome activation, ZGA) (25). These changes are prerequisite for acquiring different cell fates and specific morphological forms during development (22, 24).

The dramatic changes that occur at the onset of MBT in aquatic animal species are precisely timed in the embryo. The nuclear to cytoplasmic ratio (N/C), which gradually increases due to the reductive nature of early divisions plays a critical role in triggering MBT (24, 26, 27). In parallel, gradual dilution of maternally expressed transcription repressors and reduction of replication and transcription factors, resulting from the lack of transcription, were shown to control ZGA (17, 24, 28–31). Whether and how these factors are coordinated between single cells in the embryo is largely unknown. Previous work by Kane and Kimmel describing the changes that occur during MBT in zebrafish embryos suggested that the timing of MBT onset is more similar in immediate linear ancestor cells (e.g. sibling cells) than in cells residing at physical proximity (25). Yet, how cells distinguish their siblings from other neighboring cells has not been defined.

Regulating abscission timing can potentially affect coordination between sibling cells versus non-sibling cells by maintaining connectivity between cells through intercellular bridges. Here, we characterized abscission timing in zebrafish embryos in relation to MBT and found that abscission is delayed until the 10^th^ division cycle. Consistent with the abscission delay, expression levels of abscission related genes, including the core ESCRT component Chmp4bb, were found to rise only at MBT. Prolonged inhibition of Chmp4bb expression via morpholino, delayed abscission beyond the 10^th^ cell cycle. Prior to MBT, sibling cells continued to divide rapidly, without resolving their intercellular bridges, giving rise to the formation of inter-connected cell clusters. Notably, interconnected sibling cells exhibited a similar pattern of MBT-associated transcription onset and cell shape changes, compared to non-connected neighboring cells in the embryo. Collectively, our data indicate that abscission is part of the MBT switch and suggest that cells can temporally coordinate specific cellular behaviors by maintaining cell connectivity through intercellular bridges, which may ultimately influence their developmental program.

## RESULTS

### Characterizing cytokinetic abscission in blastula stage

We first set to describe the spatiotemporal behavior of abscission during the blastula stage in zebrafish embryos. Formation of the intercellular bridge is characterized by compaction of the mitotic spindle microtubules. Proteins that accumulated at the cell equator during anaphase, such as CEP55, condense during this process, giving rise to the formation of a dense structure called the midbody (3, 32). Cleavage of the intercellular bridge is characterized by narrowing of the microtubule stalk at either side of the midbody, followed by the release of a midbody remnant (3, 4, 33). To visualize these features in developing embryos, single cell embryos were injected with fluorescently labeled tubulin and with an mRNA encoding for the midbody protein Cep55 tagged with mCherry. This procedure did not affect the ability of the embryo to develop into a fully differentiated fish, indicating that the developmental program was not perturbed by our manipulation (Fig. S1*A*). Injected embryos were then detached from their chorions, placed in specially designed agar chambers, previously described by Mitchison and colleagues, and imaged live using a spinning disk microscope (Fig. 1A, Fig. S1*B*) (34). Embryos imaged under these conditions divided normally under the microscope and followed the previously described timeline for zebrafish at the blastula stage (Fig. 1*B*) (25). Importantly, using this setup, mitotic divisions and intercellular bridges could be readily recorded over time in developing embryos at blastula stage (Fig. 1*B*, *C* Movie S1, Movie S2).

**Figure 1:**
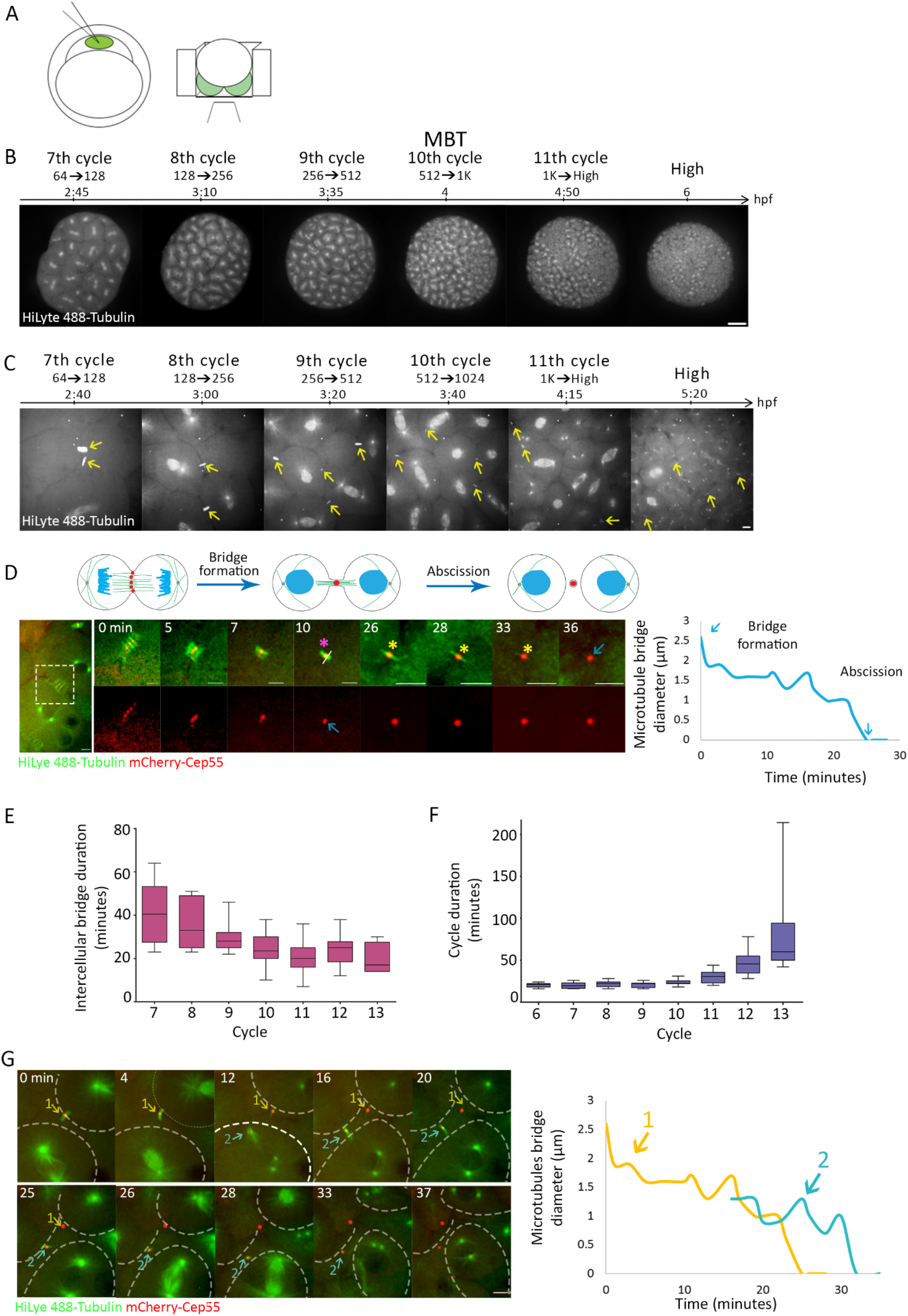
Cytokinetic abscission in zebrafish embryos at blastula stage. (***A***) Schematic illustration of the experimental setup used to visualize abscission in live zebrafish embryos during blastula. One-cell-stage embryos were injected with mRNA encoding for specific proteins and with HiLyte 488-Tubulin (left panel), dechorionized and mounted in specialized custom-made agar chambers (described in methods and Fig. S1) with the animal pole facing the cover slip, and imaged using an inverted spinning disk (right panel). (***B***) Whole embryos were visualized throughout blastula stage in live embryos. Shown are selected frames from the time lapse movie demonstrating mitosis at 7^th^ to 11^th^ cell divisions using Tubulin labeling. Maximum intensity projection of 3D volumes (30 Z slices at 1 μm) taken from a representative zebrafish embryo are shown (Movie S1). Scale bar 100 μm. (***C***) Visualizing intercellular bridges in live embryos at blastula stage. Embryos were imaged as in B, but with higher magnification for visualization of intercellular bridges. Maximum intensity projections of 3D volumes (30 Z slices at 1 μm) taken from selected timepoints of a representative developing zebrafish embryo video recording are shown (Movie S2). Intercellular bridges are marked with yellow arrows. Scale bar 10 μm. (***D***) Measuring intercellular bridge duration in zebrafish embryos. Schematic illustration of the major steps in cytokinesis that are visualized in the movie sequence are shown in upper panel. Sequential time frames from a live imaging recordings of a 512-cell stage embryo injected with HiLyte488-Tubulin (green) and mRNA encoding to mCherry-cep55 (red) are shown in bottom panel (Movie S3). A zoomed-out image of the two dividing cells is shown to the left. The midbody is marked with astrisks. The first time-point in which a single microtubule stalk with a packed Cep55 puncta were observed (magenta asterisk) was defined as the time of bridge formation. Plot on the right: the diameter of the microtubule stalk was measured at the rim of the midbody (white line), was plotted through time and was used for determining the duration of the intercellular bridge. Zero diameter indicate complete severing and represent the time of abscission. This analysis was repeated for each of the intercellular bridges measured in this study. Blue arrows represent the bridge formation to abscission. Scale bar 10 μm. (***E***) Averaged durations of intercellular bridges (measured as described in D) formed at different cell cycles (data was obtained from 5 embryos and a total of 49 intercellular bridges) All error bars are SD. (***F***) Averaged cell cycle duration measured in embryos at different division cycle. Values were measured for individual cells (as described in methods) and the averaged values obtained for each embryo were plotted according to the division cycle (N=9 embryos). All error bars are SD. (***G***) Live imaging of dividing cells in zebrafish embryos at 8^th^-10^th^ cell cycles. Maximum intensity projection of 3D volumes (30 Z slices at 1 μm) taken from selected time-points of a representative developing zebrafish embryo video recording, are shown (Movie S4). Microtubules, green; Cep55, red; intercellular bridges, arrows. Formation of spindles (green) indicate mitosis. Cells outlines are marked with dashed lines. The durations of two intercellular bridges observed in the movie sequence was (as described in *D*) are plotted to the right. Colors in plot correspond to arrows in the movie sequence. Note that the first intercellular bridge (yellow) formed at the end of the previous cycle and was not resolved when the second bridge (blue) formed at the end of the following mitotic division. Scale bar 10 μm.

Overall bridge formation and abscission followed the previously described characteristics of CEP55 condensation, bridge narrowing and midbody release, allowing us to measure the duration of intercellular bridges formed at different division cycles (Fig. 1*D*, Movie S3) (2, 32, 33, 35, 36). We found that intercellular bridges formed at subsequent division cycles persisted for shorter times, ranging from over 40 minutes in 7^th^ cell cycle to ~20 minutes in 11^th^ and 12^th^ cell cycles (Fig. 1*E*). This was in contrast to the overall cell cycle duration, which exhibited the previously characterized fast divisions until the 10^th^ cell cycle (every ~20 minutes) and was followed by gradual elongation of the cell cycle duration, to over an hour (Fig 1*F*) (23, 25). The time duration of intercellular bridges (~ 40 minutes) relative to the short division times (~ 20 minutes) in 7^th^ and 8^th^ cell cycles indicates that intercellular bridges formed in these cycles continue to exist in the following cycles (9^th^ and 10^th^). Indeed, live imaging of embryos at these stages revealed that cells connected via intercellular bridges continued to the next mitotic cycle ultimately forming new intercellular bridges without resolving the previous ones (Fig. 1*G*, Movie S4). Thus, prior to MBT, individual cells can be connected by intercellular bridges to more than one sibling cell.

### Abscission is inhibited prior to MBT

As we found that cells can be connected by multiple intercellular bridges before MBT, we wondered whether intercellular bridge duration and abscission timing are related to the MBT switch. To this end, we traced individual bridges over time in single embryos and measured the precise timing of abscission in the developing embryo. Interestingly, we could not detect any abscission event prior to MBT (Fig. 2*A*). At post MBT cycles, abscission of bridges that formed either before or after MBT was observed. These results explain the prolonged durations observed for intercellular bridges that formed in early cell cycles (6^th^ to 9^th^) and suggests that abscission is inhibited prior to MBT.

**Figure 2.**
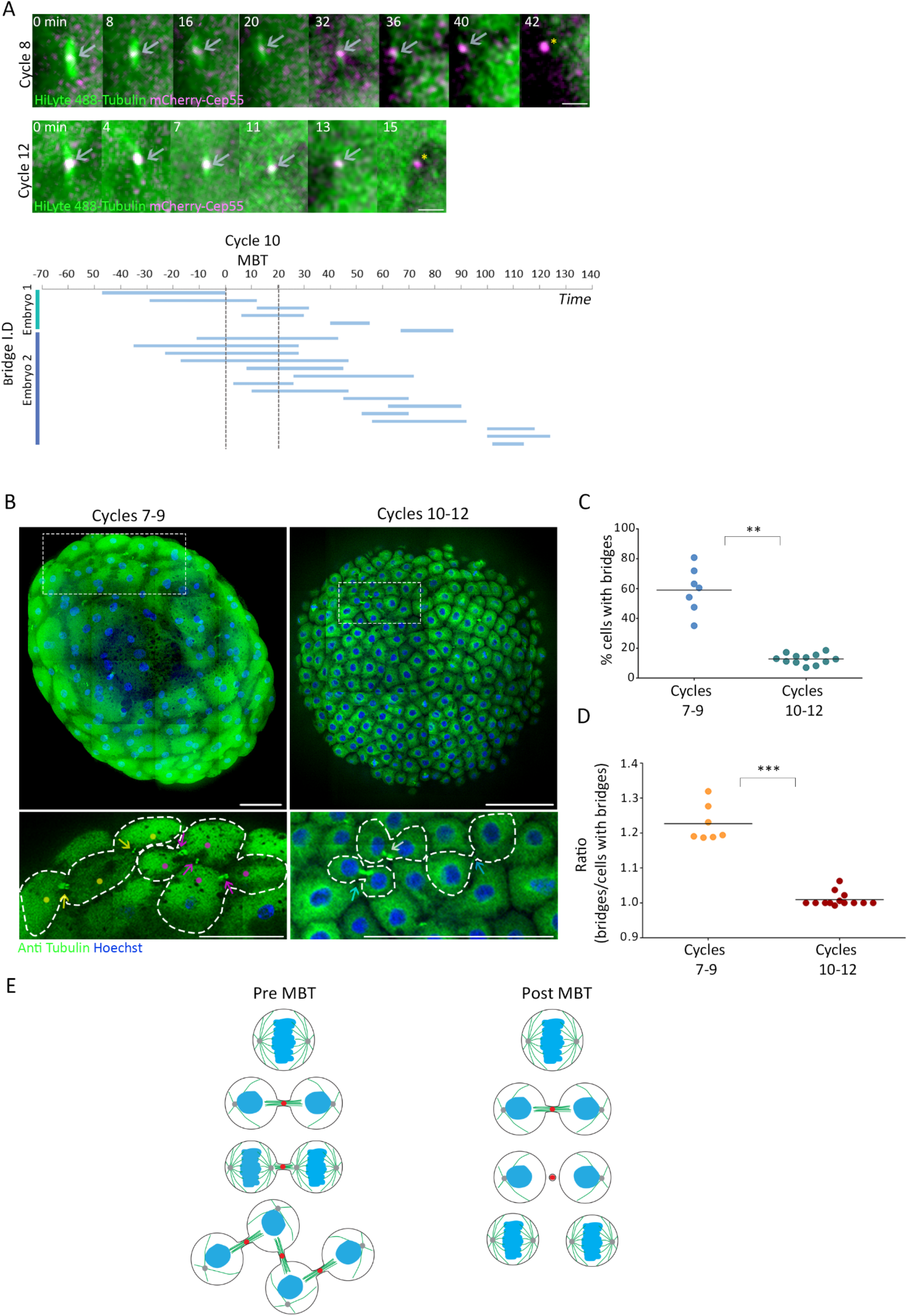
Abscission is delayed prior to MBT. (***A***) Embryos injected with mRNA encoding to mCherry-Cep55 (magenta) and HiLyte488-Tubulin (green) were imaged live in 3D and individual intercellular bridges were track and measured over time (as described in Fig 1*D*). Upper panels: maximum intensity projection images (10 Z slices of 0.7μm) of representative intercellular bridges formed pre- (8^th^ cell cycle) or post-MBT (12^th^ cell cycle), as indicated. Gray arrows, intercellular bridges; Yellow astrisks, abscission completion. Scale bar 5 μm. Bottom panel: timeline of intercellular bridges obtained from single embryos in relation to MBT. Time 0 indicate the entrance to the 10^th^ cell cycle (MBT onset). Each line represents an individual intercellular bridge (from formation to abscission). Shown are intercellular bridges from two representative embryos. (***B***) Tile images of whole embryos fixed before (7^th^-9^th^ cycles, left panel) or after MBT (10^th^-12^th^ cycles, right panel) and stained with anti-Tubulin and Hoechst. Enlarged images of the regions in dashed rectangles in upper panels are shown on the bottom. Dots indicate cells; arrows indicate intercellular bridges; dashed lines re used to mark the cells outlines. Clusters of connected cells are labeled with dots and arrows with similar colors. Scale bar 100 μm. (***C***) Percentage of cells with intercellular bridges visualized in embryos fixed at 7^th^-9^th^ cell cycle (average = 59% ± 12.21%, n=7 embryos) and in 10^th^-12^th^ cell cycle (average = 12.8%±3.43%, n=12 embryos). The percentage of cells in late cytokinesis in pre-MBT embryos (7^th^-9^th^ cell cycle) was significantly higher than in post-MBT embryos (10^th^-12^th^ cell cycles) according to *t*-test (*t*_0.05,6_=7.921, *p*=0.0002). (***D***) The ratio of intercellular bridges per cell at late cytokinesis was measured in pre- and post-MBT embryos, as described in methods, and was plotted on the graph. Averaged ratios for pre-MBT embryos = 1.009±0.019 (N=7 embryos); post MBT embryos = 1.22±0.048 (n=12). Ratios in pre-MBT embryos are significantly higher than in post-MBT embryos (Mann-Whitney U test, *U*_7,9_=0 *p*=0.0002). (***E***) Schematic model of intercellular bridges connection in pre- and post-MBT embryos. Prior to MBT, cell division leads to the formation of two sibling cells that are connected by a single intercellular bridge. Before abscission is completed, these cells enter the next division cycle and new intercellular bridges are formed. As a result, a cluster of 4 cells that are inter-connected by 3 bridges is formed. Post-MBT, cell division is followed by formation of a single intercellular bridge that is resolved prior to the next cell division cycle and, hence, inter-connected clusters are not formed.

The findings that abscission does not occur during the rapid mitotic cycles prior to MBT suggests that cells at early blastula accumulate intercellular bridges over time. To estimate the extent of this phenomenon, intercellular bridges were visualized in whole embryos that were fixed at either early or late blastula stages (pre-MBT, 7^th^-9^th^ cell cycles; post-MBT, cycles 10^th^-11^th^ cell cycles) and the number of bridges was compared across stages (Fig. 2*B*). As expected by the high rate of synchronized divisions prior to MBT, the percentage of cells connected by intercellular bridges was considerably higher in pre-MBT compared to post-MBT embryos (60% and 10% respectively) (Fig. 2*C*). Interestingly, while in post-MBT embryos, cytokinetic cells exhibited the typical morphology of a pair of sibling cells connected to each other with an intercellular bridge, in pre-MBT embryos, cytokinetic cells often obtained intercellular bridge connections to more than one cell, forming local clusters of inter-connected sibling cells (Fig. 2*B*). To quantify this effect, we measured the number of bridges connected per cell in pre- and post-MBT embryos. In post-MBT embryos (10^th^-12^th^ cell cycles), we observed the typical 1:1 ratio, indicating that each cell is connected by a single bridge. However, in pre-MBT embryos (7^th^-9^th^ cell cycles), we observed a significantly higher ratio of 1:1.24, indicating that on average each cell is connected by more than one intercellular bridge (Fig. 2*B,D*).

Next, we aimed to validate that the persisted intercellular bridges observed in pre-MBT embryos are functional and allow the exchange of cytosolic components between cells, as previously demonstrated in other cellular contexts (10–13). To this end, we injected fluorescently labelled dextran into a single cell at a 128-cells (7^th^ cycle) live embryo and tracked its diffusion over time. Dextran quickly diffused between inter-connected cells in the same cluster, but was excluded from other neighboring cells (Fig. S2, Movie S5). Collectively, these data suggest that prior to MBT cells continue to divide rapidly without resolving their intercellular bridges, which allow the formation of clusters of interconnected cells that can exchange cytosolic material (Fig. 2*E*).

### The spatiotemporal expression of ESCRTs is involved in cytokinetic abscission at blastula stage

MBT is manifested by the onset of massive transcription and global gene expression. While proteins that are essential for the first division cycles are maternally expressed, other proteins begin to transcribe from maternally deposited mRNA at early blastula stage (minor wave) and most proteins are expressed after MBT onset (major wave) (24, 37). The temporal inhibition of gene expression prior to MBT raises the possibility that expression absence of abscission relate genes is associated with the observed abscission delay. To test this notion, we analyzed the temporal transcription of 48 abscission related genes across eight developmental stages, before and after MBT, using the gene expression atlas of zebrafish embryogenesis (38) (Fig. S3*A*). Among the 48 abscission genes, we focused our analysis on the 17 genes in the zebrafish genome that belonged to the ESCRT machinery. The ESCRT machinery is a multiprotein complex that actively execute scission of the intercellular bridge and is composed early components that facilitate membrane recruitment (CEP55, TSG101, ALIX) and late components (CHMP1-7, IST1 and VPS4) that drive membrane fission (5, 39) (Fig. 3*A*). Therefore, the availability of ESCRT components and particularly of late ESCRT components can directly affect abscission timing.

**Figure 3:**
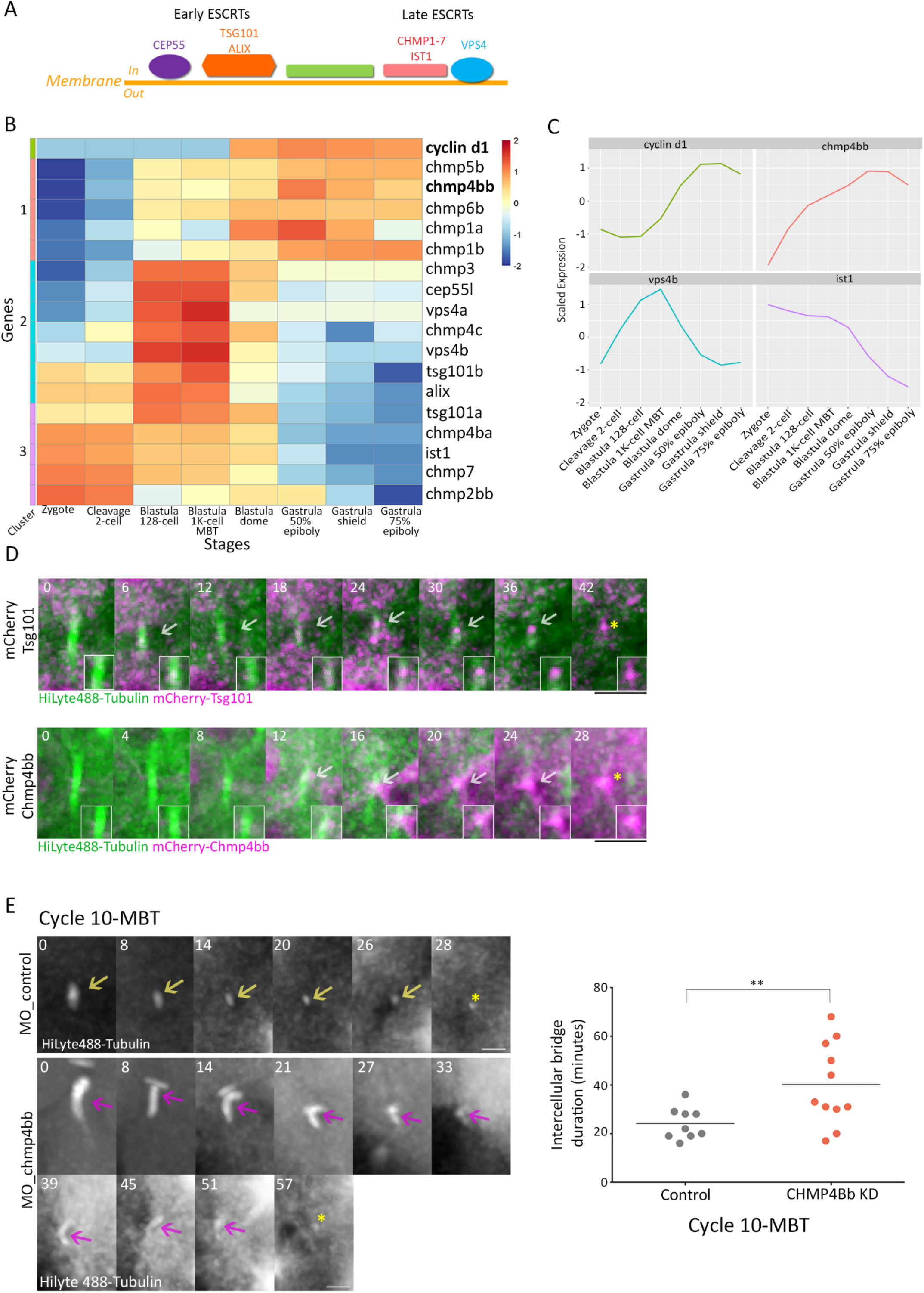
ESCRT proteins are involved in abscission during zebrafish embryogenesis. (***A***) Schematic representation of the canonical ESCRT cascade. Early ESCRT proteins such as CEP55, ALIX and TSG101 recruit late ESCRTs (CHMP1-7 and IST1) and VPS4 to the inner side of the plasma membrane, where scission occurs. (***B***) Heatmap clustering of mRNA of ESCRT genes. RNA sequencing data from eight developmental stages was downloaded from White et. al. (38). Hierarchical clustering of the expression levels of the zygotically activated gene *cyclin d1* and 17 ESCRT genes across the 8 developmental stages are presented. Rows represent individual genes and columns represents developmental stages. Colors in heatmap indicate the expression levels of each gene across the samples, relative to its mean expression and trimmed to range [-2, 2] (see Fig. S3*B* for raw data). Expression patterns were divided into 3 clusters (Cluster 1-3, color coded left to heatmap). Genes within clusters were sorted according to their scaled expression in the Zygote stage. (***C***) Normalized mRNA levels obtained for *cyclin d1* and representative ESCRT genes plotted through time. X axis, developmental stages; Y axis, normalized expression levels. (***D***) Visualizing ESCRT proteins in cytokinetic cells of zebrafish embryos at blastula stage. Embryos were injected with mRNA that encodes for Tsg101 or Chmp4bb fused to mCherry (magenta) and with HiLyte 488-Tubulin (green). Shown are maximum intensity projection images (10-25 Z slices of 0.7 μm intervals) take from a time lapse movies of representative cells imaged in embryos at 12^th^ cell cycle. Top panel: mCh-Tsg101, data was reproduced in 2 embryos and 14 intercellular bridges. Bottom panel: Chmp4bb, Data was reproduced in 6 embryos and 20 intercellular bridges. Both proteins localized to the intercellular bridge (gray arrow) and were detected in the midbody remnants after abscission (yellow asterisk). Time 0 (top left in each frame) indicate the time of bridge formation. Scale bar 10 μm. Zoomed-in images of the midbody are shown as insets in each frame. (***E***) Depletion of Chmp4bb causes abscission delays in embryos at MBT. One-cell-stage Z zebrafish embryos were injected with morpholino oligos (control or Chmp4bb specific) and intercellular bridges were recorded through time in embryos at 10^th^ cell cycle. Shown are maximum intensity projection images (6-10 Z slices of 1 μm interval) taken from movies of representative intercellular bridges (movie S6). Asterisks represents abscission completion. Scale bar 5 μm. Right panel: Intercellular bridge durations measured for individual intercellular bridges (as described in Fig. 1*D*) in control and knock down (KD) embryos. Averaged bridge duration in Chmp4bb KD embryos = 40.09±16.82 (n=11 bridges); in control embryos = 24.11±6.47minutes (n=9 bridges). Intercellular bridge duration was significantly longer in Chmp4bb KD embryos compared to control embryos (unpaired *t*-test with Welch’s correction for unequal variances *t*_0.05,13_=2.9, *p*=0.012).

Using cluster analysis of the temporal transcription of different genes, three different expression patterns were found for ESCRT genes (clusters 1-3). The expression pattern of genes in cluster 1 was characterized by low expression prior to MBT that elevated afterward, resembling the expression pattern of *cyclin d1*, which was used as a reference for zygotic genes (40, 41). The expression pattern of genes in cluster 2 was characterized by elevated expression at 128 cells stage correlating with the expression levels previously documented for minor wave genes (37). Genes in cluster 3 showed relatively high expression levels from 1 cell stage (zygote) and can therefore be considered as maternally expressed genes. Among the 17 ESCRT genes analyzed, five genes including *tsg101a* and *ist1*, were found in cluster 3, suggesting that these genes are maternally expressed. Seven ESCRT genes including *cep55, vps4a and vps4b* were in cluster 2 and another five genes including *chmp4bb* were in cluster 1, indicating that more than half of the ESCRT genes are likely not being deposited maternally (Fig. 3*B,C*, Fig S3*B*).

As CHMP4B and VPS4 manifest the minimal evolutionary conserved complex required for membrane scission (42), we further analyzed their protein expression levels. Using western blot analysis, we found that Vps4 could be detected at embryos of 256 cells while Chmp4bb could only be detected at MBT or later, supporting the spatiotemporal expression patterns of these genes and indicating, in particular, that *chmp4bb* is not expressed prior to MBT (Fig. S3*C*). Live imaging of fluorescently tagged versions of early ESCRTs (Tsg101) and late ESCRTs (Chmp4bb and Vps4) showed that they all localize to the intercellular bridge prior to abscission in cells at late blastula stage, confirming that ESCRTs are involved in cytokinetic abscission of cells in early zebrafish embryos (Fig. 3*D*, Fig. S4*A*). The additional Chmp4B homolog in the zebrafish genome, Chmp4ba, that has a similar sequence homology to the mammalian CHMP4B, could not be detected at intercellular bridges in the embryo. We therefore concluded that, at least during embryogenesis, Chmp4bb is the functional CHMP4B homolog in zebrafish cytokinetic abscission (Fig. S4*B,C*). Altogether, these results suggest that ESCRTs are involved in abscission of intercellular bridges of embryos at blastula stage and that the availability of specific ESCRT components including Chmp4bb can potentially be involved in the abscission delay observed at these at these embryonic stages.

To test the effect of ESCRT expression on abscission timing in cells at blastula stage, we used a *chmp4bb* morpholino, which reduced Chmp4bb levels by 70% (compared to embryos injected with a control morpholino sequence) (Fig. S4*D*). Live imaging of cytokinetic cells in embryos injected with *chmp4bb* morpholino revealed a significant abscission delay at 10^th^ cell cycle (40.1±16.8 minutes vs 24.1±6.47 minutes) (Fig. *3E*, Fig. S4*E*, Movies S6*A*, S6*B*). No major difference in intercellular bridge durations were recorded in cells at the 9^th^ cell cycle, in accordance with the lack of Chmp4bb expression at these stages (Fig. S3*C*). A noticeable, but milder delay was observed at the 11^th^ cell cycle, and no delay was observed at later cycles. These results indicate that depleting Chmp4bb levels, in early embryos, considerably inhibit abscission at MBT while only mildly affecting later stages. It is possible that high levels of Chmp4bb are specifically crucial during MBT to allow the severing of all the intercellular bridges that accumulated in the embryo. Together with the zygotic expression pattern of *chmp4bb*, these results suggest that the levels of Chmp4bb play a role in timing bridge cleavage in the embryo.

### Similar transcription pattern and cell shape changes in inter-connected cells at MBT

Next, we wondered what could be the developmental role of stalling abscission prior to MBT. As the global transcription of the zygote genome mainly occurs at the MBT, we set to examine the onset of transcription in relation to cell-cell connectivity in zebrafish embryogenesis. Initially, we characterized the overall transcription levels in single cells of pre- and post-MBT embryos. To this end, one-cell-stage embryos were injected with a modified uridine, that is capable of binding a fluorescent dye via a click-reaction (Fig. 4*A*). Upon transcription in the embryos, the modified uridine is incorporated into the newly synthesized transcripts and can serve as a reporter for the overall transcription level in the nucleus, as previously described (43). Using this analysis, we identified four transcription patterns in embryos at blastula stage: 1) No transcription, only background uridine labeling. 2) low transcription, uridine labeling at two foci in the nucleolus. 3) high transcription, the entire nucleus is labeled with uridine. 4) dividing cells, no uridine labeling accompanied by condensed chromatin staining at the cell equator (Fig. 4*B*). These transcription patterns were observed different embryonic stages: in early stages (7^th^ cell cycle, 128 cells), no transcription was observed, in the 9^th^ cell cycle (512 cells), cells exhibited the characteristic staining of low transcription, and at later stages (12^th^-15^th^ cycles, high and dome) transcription levels were high (Fig. 4*C*, Fig. S5*A*).

**Figure 4:**
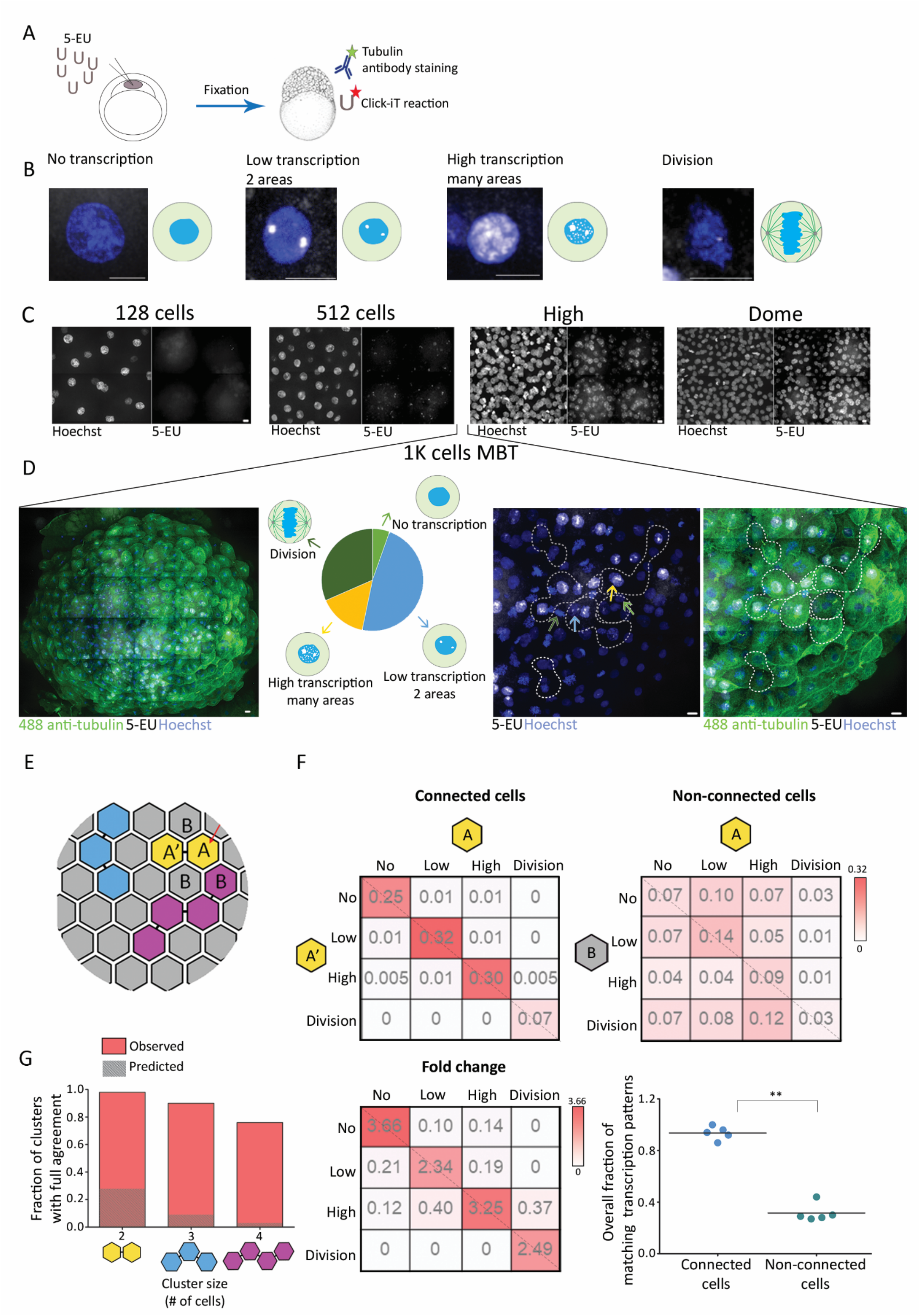
Cells connected by intercellular bridges exhibit similar transcription pattern. (***A***) Schematic illustration of the experimental setup. One-cell-stage embryos were injected with 5-ethynyl uridine (5-EU) and fixed at the indicated stages. Embryos were stained with anti-Tubulin (for intercellular bridge detection), Hoechst (for nuclear detection) and click-labeled using Alexa594-azide for detection of mRNA incorporated 5-EU. (***B-D***) Transcription patterns observed in zebrafish embryos at blastula stage. Embryos were treated as described in A and imaged using a spinning disk confocal. (***B***) Single cells labeled with 5-EU (white) and Hoechst (blue), exhibiting the 5-EU labeling patterns observed in the embryos. Transcription patterns were classified into four different stages, based on 5-EU labeling: no, low, high and dividing cells (43). A cartoon demonstrating the transcription pattern is shown to the right of each image. Scale bar, 10 μm. (***C***) Tile images of maximum intensity projection images (15-30 z slices of 1-1.5 μm interval) of representative regions in embryos at the indicated developmental stages. Embryos at 128 cells stage (7^th^ cell cycle), no 5-EU labeling (n = 4 embryos); Embryos at 512 cells stage (9^th^ cell cycle), 5-EU in two foci (n = 4 embryos); Embryos at high and dome stages (13^th^ and 14^th^ cell ycles, respectively), dense 5-EU labeling in nucleus (n (high) = 4 embryos, n (dome) = 5 embryos). Scale bar 10 μm. (***D***) Heterogeneous transcription patterns observed in embryos at MBT (1K-cell stage, cycle 10). Left panel: A tile image of a representative embryo in cycle 10 labeled with Tubulin (green), Hoechst (blue) and 5-EU (white). Second panel: the relative distribution of the four transcription patterns (classified in B) observed in the representative embryo (no transcription, 5%; low transcription, 48%, high transcription 15%, dividing cells 32%). Note that all four transcription patterns were observed in a single embryo at MBT. Third and fourth panels: Enlarged images of subset regions in the embryo that exhibited high levels of cell-cell variations in transcription patterns. Different transcription patterns are marked in arrows at different colors (corresponding to arrows in second panel). Clusters of inter-connected cells (marked in dashed lines) were identified based on Tubulin staining and were used in the analysis described in panels E-G. Scale bar 10 μm. (***E-G***) Quantitative analysis of the similarity in transcription patterns observed between connected and non-connected cells. (***E***) Schematic representation of an embryo at MBT. Connected cells are filled with similar colors; letters, correspond to the matrixes presented in F. (***F***) Resemblance matrix of transcription patterns of embryos at MBT. Top panels: pairs of connected (left) and non-connected cells (right) (A, A’, and A, B respectively) were scored according to their transcription pattern and the probability to obtain specific combinations of transcription patterns between cell pairs was plotted in the matrix. The diagonal (dashed line) represents identical transcription patterns between cells. Color indicate probability. Data was obtained from 5 embryos (206 connected cells and 768 non-connected). Bottom left panel: fold change increase in connected cells versus non-connected cells was calculated for each combination of transcription patterns and plotted in the matrix. Bottom right panel: diagonal of resemblance. The probabilities obtained across the diagonal in each embryo were summed up and plotted in the graph. Connected cells obtained a significantly higher averaged value compared to non-connected cells (0.94±0.05 and 0.32±0.07, respectively). The two groups are significantly different according to Mann-Whitney U test (*U*_5,5_=0.8, *p*=0.003). (***G***) Clusters of 2, 3 and 4 connected cells were scored for matched transcription patterns such that only clusters in which all cells exhibited a similar transcription patterns were considered as clusters with intra-cluster agreement (red bars). Values were compared to simulated data, generated based on marginal distribution of the different transcription patterns observed in embryos at MBT (dashed bars). Notably, cells residing in inter-connected clusters exhibited considerably higher levels of agreement in their transcription patterns compared to the predicted values. 2 cells clusters: 98.6% identical clusters vs 28% in simulation (n=150 clusters); 3 cells clusters: 90% identical clusters vs 9.1% in simulation (n=30 clusters); 4 cells clusters: 76.9% identical clusters vs 3.1% in simulation (n=12 clusters).

Interestingly, while in embryos at early and late blastula stage (corresponds to 128-512 cells and high-dome, respectively) a uniform transcription pattern was observed throughout the embryo, considerable variations in transcription patterns were observed between individual cells in embryos at the 10^th^ cell cycle (Fig. 4*D*). In fact, all four transcription patterns could be observed in a single embryo at this stage. This heterogeneity in transcription levels during MBT, indicate that cells autonomously regulate the timing of their transcriptional onset, as previously suggested (25).

Given that the MBT switch is thought to be driven by the ratios of cellular components and that diffusion occurs through intercellular bridges of inter-connected sibling cells, we asked whether inter-connected cells exhibit similar transcription patterns. For that, embryos at the 10^th^ cycle were labeled both for transcription (using modified uridine) and for intercellular bridges (using anti-tubulin antibodies) and the transcription patterns of pairs of cells connected by bridges (connected cells) and adjacent pairs of neighboring cells that are not connected by a bridge (non-connected cells) was determined (Fig. 4*E*, Fig. 4*F*, A-A’ versus A-B correspondingly). We then pooled all connected and non-connected cell pairs and calculated the *transcription agreement probability matrix*, where the bin at row *r* and column *c* holds the probability for a pair (connected or non-connected) to have one cell in the transcriptional pattern corresponding to *r* and the other in the pattern corresponding to *c* (Fig. 4F). The accumulated probabilities at the diagonal of the agreement matrix is the probability of two cells within a pair to have the same transcriptional pattern. This analysis revealed that connected cells exhibit higher degree of agreement in their transcription pattern compared to non-connected neighboring cells (Fig. 4*F*, 0.94 versus 0.33 correspondingly). This observation of matched transcription patterns was not limited to pairs of cells. In fact, the intra-cluster transcriptional agreement was even more striking in clusters of three and four connected cells in respect to a null model that assumed no spatial relations (Fig. 4*G*, Methods). These data establish that inter-connected cells are subjected to similar regulation of transcription.

Last, we asked whether cellular properties downstream of transcription are coupled in inter-connected cells at MBT. We focused on changes in cell polarization and shape, that were previously described to occur during the MBT switch (25). We thus compared the deviation in the cell’s aspect ratio (i.e. the major axis of each cells divided by its minor axis) in pairs of adjacent connected cells versus non-connected neighboring cells (Fig. S5*B*, *Methods*). More specifically, we considered triplet of neighboring cells, that included two connected cells and one non-connected cell and measured their aspect ratio (Fig. S5*B*. Movie S7). We then calculated, for each triplet, the absolute difference in the aspect ratio between two connected and non-connected cells, as a measure of morphological similarity. The difference in the aspect ratio of connected pairs was significantly smaller compared to neighboring non-connected pairs, as verified by unmatched (0.23±0.05 compared to 0.35±0.19, respectively) or matched analysis - directly comparing cell pairs from the same triplet (Fig. S5*C,D*). These results establish a stronger agreement of cell shape in connected cells. Taken together with the similarity in transcription pattern, these results suggest that abscission is an integral part of MBT that allows coordinating cellular behavior during the MBT period.

## DISCUSSION

In this work, we showed that cells do not resolve their intercellular bridges prior to MBT and that abscission commence at the 10^th^ division cycle. This phenomenon was associated with the formation of clusters of cells that are connected by intercellular bridges and exhibit matching transcriptional and morphological features during MBT, suggesting that cells can communicate through their intercellular bridges. Indeed, intercellular bridges were previously shown to enable the exchange of molecules between daughter cells in both cultured mammalian cells and in whole organisms (*Drosophila* and *C. elegans*). Additionally, protein exchange was documented between cells in zebrafish embryos at gastrula stage, that maintained long ranged intercellular bridges (13). Consistently, we showed that clusters of inter-connected cells exchange cytoplasmic dextran. Together, these findings indicate that delayed abscission prior to MBT plays a role in the formation of inter-connected cell clusters in the embryos that coordinate their behavior at early stages of embryogenesis.

In 1995, Kimmel and colleagues provided a detailed description of the stages of embryonic development in zebrafish (23). In this seminal work, they report that in the first four cell divisions of zebrafish embryos furrowing is not complete and the cells remain partly connected to one another and to the yolk. Once the embryo reaches the 32-cell stage, furrowing is complete and the cells are fully separated from one another. However, several lines of evidence suggest that cells at these stages (32 −1K cells) sustain memory of their lineage. For example, in 1995 Kane and Kimmel reported that although cells autonomously set on the MBT switch, sibling cells exhibit more similar behaviors compared to other non-connected neighboring cells (25, 44). Additionally, in cell lineage reconstruction experiments, cells in embryos at 128-512 cells stage appeared to reside in clusters based on their 8 cells blastomere lineage, raising the possibility that these cells are physically connected to one another in some fashion (44). More recently, the levels of zygotically activated genes, were shown to be more similar in daughter cells compare to other neighboring cells in embryos at blastula stage (29). All these observations indicated that sibling cells in the embryo maintain a cellular memory of their lineage, however, how this memory is transmitted between the cells was not addressed. Given our observations of persistent intercellular bridges prior to MBT, we propose that the memory observed between cells at these stages originate from their cell-cell connectivity through intercellular bridges. Thus, we would like to suggest the following scenario for zebrafish development: early embryogenesis begins by incomplete furrowing during mitosis until the embryo reach 32-cells stage. Then, complete furrowing occurs but abscission does not commence and cells remained connected to one another until they reach 1K-cells stage (MBT). After MBT, complete furrowing followed by abscission give rise to completely independent cells.

What governs the onset of abscission at MBT is still not fully understood. Our data suggests that the ESCRT-III component Chmp4bb may contribute to this process. Notably, levels of the CHMP4B homolog in *Drosophila*, Shrub, were also shown to regulate abscission timing in the *Drosophila* germline and loss of Shrub induced the formation of egg chambers with 32 connected cells (10). Given the fundamental role of CHMP4B in executing ESCRT mediated fission *in vitro* and *in vivo*, it is possible that the cellular levels of Chmp4bb are tightly regulated during development and are served to control cell-cell connectivity in the embryo by regulating abscission timing (10, 33, 36, 45, 46). Expression levels of other abscission related genes, such as *rhoab, rab11a, arf6b* and additional late ESCRT genes such as *CHMP3* and *CHMP1B*, were found to increase toward the MBT, implying that additional genes may regulate abscission timing in embryo to ensure precise developmental program (Fig. S3*A* and 3*B*). Regardless of the mechanism, abscission should be regarded as part of the MBT switch and cell-cell connectivity should be considered when addressing collective cellular behavior in early zebrafish embryos.

## METHODS

### Fish maintenance

The animal work performed in this study was approved by the Ben Gurion University Institutional Animal Care and Use Committee (BGU520916). AB wild type zebrafish were maintained and bred at 28 °C according to standard protocols.

### mRNA and morpholino

Cep55l, Tsg101, Chmp4bb, Chmp4ba zebrafish (*Danio rerio*) coding sequences were amplified by PCR from zebrafish cDNA using specific primers (Table S1) and subcloned into mCherry-C1 vector (Clontech, Mountain View, CA). hVPS4A in pEGFP-C1 was previously described in Elia et. al. (33). All plasmids then subcloned into pCS2+ vector (kindly provided by Gil Levkowitz, Weizmann Institute of Science, Israel). mRNA encoding for mCherry-cep55 mCherry-tsg101, mCherry-chmp4bb, mCherry-chmp4ba were generated by *in-vitro* transcribed using mMessage mMachine SP6 kit (AM-1340 Invitrogen) and purified using RNeasy mini kit (74104 Qiagen). Morpholino oligos against chmp4bb and standard control oligos (Gene Tools, Inc.) were designed and managed according to manufacture protocol (Table S1). Two morpholino oligos were injected and analyzed by western blot analysis for chmp4bb knock down (MO_1chmp4bb and MO_2chmp4bb) (Fig. S4*D*). MO_1chmp4bb was found to be more efficient by western blot analysis, and was therefore used in all the presented experiments.

### Embryos microinjection and mounting for live imaging

mRNA (0.8ng-1ng) or morpholino oligos (5ng) were microinjected into wild type embryos at one-cell stage together with 1.3 ng HiLyte Fluor™ 488-Tubulin (TL488M, Cytoskeleton, Inc.) or 1.3 ng TAMRA Rhodamine Tubulin (TL590M, Cytoskeleton, Inc.) and 0.1% phenol red (P0290, Sigma-Aldrich) using microinjector (World Precision Instrument, Sarasota, FL) and kept in Danieau buffer (DBx1) (174 mM NaCl, 21 mM KCl, 12 mM MgSO4, 18 mM Ca(NO3)2, 15 mM HEPES). Under these conditions, embryos were viable and showed normal cell division and physiological morphology at blastula stage (Fig. 1*B*, Fig. S1*A*). Injected embryos were dechorionized using fine tweezers and placed in specialized agar chambers for imaging.

Preparation of agar chambers: specialized agar wells designed for mounting zebrafish embryos at blastula stage for live imaging were prepared according to Whur et. al (34). In short, a mold with 8 wells, each 0.6-0.7 μm^2^, was placed in a 35 mm glass bottom μ-dish (81156, ibidi) (Fig S1*B*). Wells were mounted by pouring 1 ml of 2.5% melted agarose (50080, Lonza Rockland, ME in Danieau buffer (DBx1) into the μ-dish with the mold inside. After agar solidifying, the mold was carefully removed and dechorionized embryos were inserted to the chamber with the animal pole facing the coverslip and covered with DBx1 (Fig 1*A*).

### Whole-mount immunostaining

Embryos at 2-2.75 hpf (7^th^-9^th^ cell cycles, 128-512 cells) and at 3.5-4.5 hpf (11^th^-12^th^ cell cycles) were fixed (4% PFA, overnight at 4°C), washed twice in PBS supplemented with 0.1% Tween20, dechorionized with fine tweezers and placed in a 24-well dish coated with 1% agar. Embryos were then permeabilized in 0.25% trypsin-EDTA (Biological industries, Israel) in Hank’s balanced salt solution (Thermo Fisher Scientific, Waltham, MA) for 10 minutes on ice, washed three times for 30 minutes in washing buffer (PBS plus 0.2% Triton X-100) and post-fixed with 4% PFA for 1 hour at room temperature. Embryos were blocked for 2 h at room temperature in blocking solution (10% fetal bovine serum, 1% BSA and 0.2% Triton X-100 in PBS) and incubated with monoclonal anti α-Tubulin (DM1A; Sigma-Aldrich, 1:1000, overnight). After washing (x3, 20 minutes) embryos were incubated with a secondary antibody Alexa Flour 488 anti-mouse (A21202, Invitrogen, 1:1000, overnight) and washed again (x3, 20 minutes). Finally, embryos were stained with Hoechst 1:1000 (H-3570 Invitrogen) for 30 minutes, washed (x2, 15 minutes) and kept in 25% glycerol in PBS. For dissection, embryos were placed on a microscope slide in 50% glycerol solution and the yolk was cut out using a sharp scalpel and tweezers. Last, embryos were mounted in fluoromount-G^®^ (SouthernBiotech, Birmingham, AL) on a coverslip with the animal pole facing the coverslip. Samples were imaged by confocal microscopy as described below.

### Imaging and image processing

Live embryos were imaged from 8-cell stage to dome stage (13^th^ −14^th^ cell cycle) on a fully incubated confocal spinning-disk microscope at 26 °C (Marianas; Intelligent Imaging, Denver, CO) using a X40 oil objective (numerical aperture 1.3) or X20 air objective (numerical aperture 0.8). Embryos were recorded on an electron-multiplying charge-coupled device camera (Evolve; Photometrics, Tucson, AZ). A total of 25-32 confocal slices of 0.7-1.0 μm were captured for each time point at 1.5-2 minutes time interval. Only the two cell layers close to the animal pole of the embryo were imaged and analyzed.

Fixed embryos were imaged with a X40 oil objective (numerical aperture 1.4) or X63 oil objective (numerical aperture 1.4). A total of 30-100 slices at 0.6-1.5 μm intervals were acquired, providing a Z range of up to 150 μm. Tile acquisition was applied for capturing the entire embryo and 3D volumes were merged to a montage using SlideBook (version 6, Intelligent Imaging, Denver, CO). To improve visualization, images were subjected to Gaussian filtering in slidebook.

### Data analysis

Cell cycle length was defined by the morphology of microtubules at the cells. Following microtubules labeling two centrosomes at prophase and a spindle at metaphase can be clearly identified. Cell cycle length was defined as the time interval between one prophase to the next in the same cell.

To measure abscission duration, intercellular bridges were annotated and tracked through time. Interellular bridge formation was defined based on the formation of condensed microtubules fibers between the cells (Tubulin channel) with a dense Cep55 puncta at the center (Fig 1*D*). Completion of abscission was determined based on the diameter of the intercellular bridge and the formation of a midbody remnant, as previously described (2). In the absence of Cep55 expression, bridge duration was measured according to Tubulin morphology alone.

Measuring bridges to cells ratio in fixed embryos. Intercellular bridges and cells wich carry intercellular bridges were manually annotated and the ratio between them was calculated for each embryo. For calculations, intercellular bridges were counted twice (once for each cell it is connected to), resulting in a 1:1 ratio between intercellular bridges and cells, under typical conditions (two cells connected via a single intercellular bridge).

### Click-iT labeling of transcription pattern

Click-iT^®^ reaction kit was used to visualize transcription levels as previously described by Chan et al. 2019 (43). One-cell-stage embryos were microinjected with 50 pmols of 5-ethynyl uridine (5-EU, E10345), and 0.1% phenol red and kept in DBx1. Depending on the desired developmental stage, injected embryos were fixed 2-5 hpf (with 3% PFA and 0.1% glutaraldehyde mixture (overnight, 4°C) and washed three times with PBSx1. Next, embryos were then dechorionized with fine tweezers and placed in a 24-well dish coated with 1% agar for immunostaining and click-it reaction.

Immunostaining for 5-EU injected embryos: embryos were permeabilized with PBS with 0.5% Triton X-100 (PBS-T) at room temperature for 30 minutes, followed by dehydration with serial dilutions of methanol in PBS-T (25%,50%,75%,100%) for 5 minutes in each dilution. Dehydrated embryos were incubated in 100% methanol (−20°C, 2 hours) and subjected to rehydration with serial methanol in PBS-T (75%,50%,25%, 0%). Embryos were then washed, blocked (0.2% Triton X-100, 1%BSA, 10% fetal bovine serum in PBS) for 2 hours in room temperature and stained with anti α-Tubulin (DM1A; Sigma-Aldrich, 1:1000, overnight) followed by secondary antibody staining (Alexa Flour 488 anti-mouse 1:1000,2 hours in room temperature).

Click-iT reaction: embryos were incubated for 1 hour at room temperature in cell reaction mix: 5 μM of Alexa Fluor^®^ 594 azide (A10270) in Click-iT reaction buffer according to manufacture protocol (Click-iT^®^ reaction buffer kit C10269, Invitrogen). Embryos were washed twice for 10 minutes in PBS solution with 2% BSA. Finally, embryos were stained with Hoechst (0.5 hour, 1:1000), dissected and mounted as described above.

### Analysis of transcription pattern similarities in connected and non-connected cells

The agreement in transcription patterns between adjacent cells was quantified using montage images of embryos labeled with 5-EU and stained with Tubulin, as described above. First, clusters of cells connected by intercellular bridges were annotated according to Tubulin staining in specific regions in the embryos that exhibited high levels of heterogeneity in cellular transcription patterns. Next, a transcription pattern was manually assigned to cells in in the selected regions according to their 5-EU staining (see Fig. 4*B*). Then the transcription agreement probability matrix was plotted for cells connected by intercellular bridges (A, A’), and for neighboring cells that are not connected by a bridge (A, B) (Fig. 4*F*). Each bin in the column (c) and row (r) holds the fraction of cell pairs where one cell had transcription pattern c and the other transcription pattern r (i.e. the accumulated values in the matrix sum up to 1). For example, in Fig. 4 the bin at the column labeled “no” and the row labeled “high” holds the fraction of cell pair with the corresponding transcription patterns. Note, that these matrices were not completely asymmetric,(i.e. the value in bin (c,r) could slightly deviated from the value in bin (r,c) because the selection of A-A’ and A-B pairs used a common “anchor” A (see illustration in Fig. 4*E*). The disagreement between the asymmetric matrix entries was small, would converge to 0 with larger N - sampling of pairs, and did not affect the interpretation of the results in any way. The accumulated diagonal bins of these transcription agreement probability matrices held the overall fraction of adjacent cell pairs with agreement in their transcriptional patterns.

To assess the transcription agreement in cell clusters (Fig. 4*G*) we implemented a simplistic simulation to calculate the expected agreement in cell clusters under the assumption of no spatial independence in transcriptional patterns. For clusters of size 2, 3, or 4, we selected 100,000 cell clusters using the observed transcriptional pattern marginal distribution (Fig. 4*D*) and recorded the fraction of homogeneous clusters – where all cells had the same transcriptional pattern. We compared these simplistic predicted probabilities to the matching probabilities recorded in experiments (Fig. 4*G*).

### Statistics

Data was analyzed for column statistics in GraphPad Prism version 5.00 for Windows (La Jolla, CA, USA). Statistical significance was determined by *t*-test, Mann-Whitney *U* test or Wilcoxon matched-pairs signed rank test (Z) as specified in figure legends.

### RNA-seq data and visualization

RNA sequencing expression data (FPKM) of zebrafish embryos was downloaded from the Expression Atlas site (https://www.ebi.ac.uk/gxa/experiments/E-ERAD-475) (38). Eight developmental stages (zygote, cleavage 2 cell, blastula 128 cell, blastula 1k cell, blastula dome, gastrula 50% epiboly, gastrula shield, gastrula 75% epiboly) were used for the analysis. Normalized FPKM values that were lower than one was replaced by one, and all values were log2 transformed. Only the set of abscission related genes that were expressed (above zero) in at least 3 of the 8 samples were considered (48 genes). Hierarchical gene clustering was performed with hclust R function (“complete” method), using the normalized log2-transformed FPKM values of the eight developmental stages. Clusters were sorted manually and genes within clusters were sorted according to their scaled expression in the Zygote stage. For heatmaps presentation, the log2-transformed FPKM expression values of the presented genes were standardized by subtraction of each gene’s mean expression, followed by division by the standard deviation across all samples. Cyclin d1, a well-established zygotic gene was used as a positive control in our analysis was clustered into a singleton cluster and is presented in the first row of the heatmaps.

## Supporting information

Movie S1. Visualizing mitosis in live zebrafish embryos

Movie S2. Visualizing intercellular bridges in live zebrafish embryos

Movie S3. Intercellular bridge duration

Movie S4. Persistence on intercellular bridges in early embryos

Movie S5. Dextran diffusion between clusters of cells inter-connected by intercellular bridges

Movie S6. Bridge duration of control embryo

Movie S6. Bridge duration of knock down embryo

Movie S7. Changes in cells shape

## AUTHOR CONTRIBUTIONS

S.A.-L and N.E initialized the project and conceptualized all experiments. S.A.-L designed and analyzed all experiments. S.A.-L, D.N and Y.J performed experiments and assisted with analysis. S.T.G.-O performed transcriptomic analysis, N.P performed dextran injection experiments. A.Z advised on quantitative data analysis and statistics and helped writing the manuscript. R.Y.B provided guidance and help with zebrafish maintenance, injections and experiments, was actively involved in experimental design throughout the project and was involved in writing the manuscript. N.E was involved in all experimental design and data analysis and wrote the manuscript.

## ACKNOWLEDMENTS

We thank all members of the Elia and Birnbaum laboratories for their help and feedback throughout the project. We also thank members of the Peyrieras lab for hosting Shai and sharing with her their knowledge and expertise. The Elia laboratory is funded by the Israeli Science Foundation (ISF) Grant no. 1323/18. The Birnbaum laboratory is funded by the Israel Science foundation (960/2016) and the research without Borders– ST BONIFACE HOSPITAL – BEN GURION UNIVERSITY. AZ was supported by the Israeli Council for Higher Education (CHE) via Data Science Research Center, Ben-Gurion University of the Negev, Israel. S.T.G.-O. was supported by Hi-Tech, Bio-Tech, and Chemo-tech fellowship and Negev fellowship of Ben-Gurion University of the Negev.

## REFERENCES

1. P. P. D’Avino, M. G. Giansanti, M. Petronczki, Cytokinesis in animal cells. Cold Spring Harb Perspect Biol 7, a015834 (2015).

2. O. Gershony et al., Measuring abscission spatiotemporal dynamics using quantitative high-resolution microscopy. Methods Cell Biol 137, 205–224 (2017).

3. R. A. Green, E. Paluch, K. Oegema, Cytokinesis in animal cells. Annu Rev Cell Dev Biol 28, 29–58 (2012).

4. N. Elia, C. Ott, J. Lippincott-Schwartz, Incisive imaging and computation for cellular mysteries: lessons from abscission. Cell 155, 1220–1231 (2013).

5. C. Addi, J. Bai, A. Echard, Actin, microtubule, septin and ESCRT filament remodeling during late steps of cytokinesis. Curr Opin Cell Biol 50, 27–34 (2018).

6. V. Nahse, L. Christ, H. Stenmark, C. Campsteijn, The Abscission Checkpoint: Making It to the Final Cut. Trends Cell Biol 27, 1–11 (2017).

7. J. Mathieu et al., Aurora B and cyclin B have opposite effects on the timing of cytokinesis abscission in Drosophila germ cells and in vertebrate somatic cells. Dev Cell 26, 250–265 (2013).

8. O. Gershony, T. Pe’er, M. Noach-Hirsh, N. Elia, A. Tzur, Cytokinetic abscission is an acute G1 event. Cell Cycle 13, 3436–3441 (2014).

9. J. Konig, E. B. Frankel, A. Audhya, T. Muller-Reichert, Membrane remodeling during embryonic abscission in Caenorhabditis elegans. J Cell Biol 216, 1277–1286 (2017).

10. N. R. Matias, J. Mathieu, J. R. Huynh, Abscission is regulated by the ESCRT-III protein shrub in Drosophila germline stem cells. PLoS Genet 11, e1004653 (2015).

11. R. A. Green et al., The midbody ring scaffolds the abscission machinery in the absence of midbody microtubules. J Cell Biol 203, 505–520 (2013).

12. P. Steigemann et al., Aurora B-mediated abscission checkpoint protects against tetraploidization. Cell 136, 473–484 (2009).

13. L. Caneparo, P. Pantazis, W. Dempsey, S. E. Fraser, Intercellular bridges in vertebrate gastrulation. PLoS One 6, e20230 (2011).

14. C. Eno, T. Gomez, D. C. Slusarski, F. Pelegri, Slow calcium waves mediate furrow microtubule reorganization and germ plasm compaction in the early zebrafish embryo. Development 145 (2018).

15. T. Yabe et al., The maternal-effect gene cellular island encodes aurora B kinase and is essential for furrow formation in the early zebrafish embryo. PLoS Genet 5, e1000518 (2009).

16. B. Feng, H. Schwarz, S. Jesuthasan, Furrow-specific endocytosis during cytokinesis of zebrafish blastomeres. Exp Cell Res 279, 14–20 (2002).

17. M. T. Lee et al., Nanog, Pou5f1 and SoxB1 activate zygotic gene expression during the maternal-to-zygotic transition. Nature 503, 360–364 (2013).

18. A. H. Eikenes et al., ALIX and ESCRT-III coordinately control cytokinetic abscission during germline stem cell division in vivo. PLoS Genet 11, e1004904 (2015).

19. Y. M. Elkouby, A. Jamieson-Lucy, M. C. Mullins, Oocyte Polarization Is Coupled to the Chromosomal Bouquet, a Conserved Polarized Nuclear Configuration in Meiosis. PLoS Biol 14, e1002335 (2016).

20. X. Bai et al., Aurora B functions at the apical surface after specialized cytokinesis during morphogenesis in C. elegans. Development 147 (2020).

21. P. Lujan et al., PRL-3 disrupts epithelial architecture by altering the post-mitotic midbody position. J Cell Sci 129, 4130–4142 (2016).

22. J. C. Siefert, E. A. Clowdus, C. L. Sansam, Cell cycle control in the early embryonic development of aquatic animal species. Comp Biochem Physiol C Toxicol Pharmacol 178, 8–15 (2015).

23. C. B. Kimmel, W. W. Ballard, S. R. Kimmel, B. Ullmann, T. F. Schilling, Stages of embryonic development of the zebrafish. Dev Dyn 203, 253–310 (1995).

24. A. R. Langley, J. C. Smith, D. L. Stemple, S. A. Harvey, New insights into the maternal to zygotic transition. Development 141, 3834–3841 (2014).

25. D. A. Kane, C. B. Kimmel, The zebrafish midblastula transition. Development 119, 447–456 (1993).

26. J. Newport, M. Kirschner, A major developmental transition in early Xenopus embryos: I. characterization and timing of cellular changes at the midblastula stage. Cell 30, 675–686 (1982).

27. M. Zhang, P. Kothari, M. A. Lampson, Spindle assembly checkpoint acquisition at the midblastula transition. PLoS One 10, e0119285 (2015).

28. C. Collart, G. E. Allen, C. R. Bradshaw, J. C. Smith, P. Zegerman, Titration of four replication factors is essential for the Xenopus laevis midblastula transition. Science 341, 893–896 (2013).

29. Y. Hadzhiev et al., A cell cycle-coordinated Polymerase II transcription compartment encompasses gene expression before global genome activation. Nat Commun 10, 691 (2019).

30. M. Zhang, P. Kothari, M. Mullins, M. A. Lampson, Regulation of zygotic genome activation and DNA damage checkpoint acquisition at the mid-blastula transition. Cell Cycle 13, 3828–3838 (2014).

31. M. Meier et al., Cohesin facilitates zygotic genome activation in zebrafish. Development 145 (2018).

32. W. M. Zhao, A. Seki, G. Fang, Cep55, a microtubule-bundling protein, associates with centralspindlin to control the midbody integrity and cell abscission during cytokinesis. Mol Biol Cell 17, 3881–3896 (2006).

33. N. Elia, R. Sougrat, T. A. Spurlin, J. H. Hurley, J. Lippincott-Schwartz, Dynamics of endosomal sorting complex required for transport (ESCRT) machinery during cytokinesis and its role in abscission. Proc Natl Acad Sci U S A 108, 4846–4851 (2011).

34. M. Wuhr, N. D. Obholzer, S. G. Megason, H. W. Detrich, 3rd, T. J. Mitchison, Live imaging of the cytoskeleton in early cleavage-stage zebrafish embryos. Methods Cell Biol 101, 1–18 (2011).

35. H. H. Lee, N. Elia, R. Ghirlando, J. Lippincott-Schwartz, J. H. Hurley, Midbody targeting of the ESCRT machinery by a noncanonical coiled coil in CEP55. Science 322, 576–580 (2008).

36. T. Wollert, C. Wunder, J. Lippincott-Schwartz, J. H. Hurley, Membrane scission by the ESCRT-III complex. Nature 458, 172–177 (2009).

37. H. Aanes et al., Zebrafish mRNA sequencing deciphers novelties in transcriptome dynamics during maternal to zygotic transition. Genome Res 21, 1328–1338 (2011).

38. R. J. White et al., A high-resolution mRNA expression time course of embryonic development in zebrafish. Elife 6 (2017).

39. J. Guizetti, D. W. Gerlich, Cytokinetic abscission in animal cells. Semin Cell Dev Biol 21, 909–916 (2010).

40. K. T. Duffy et al., Coordinate control of cell cycle regulatory genes in zebrafish development tested by cyclin D1 knockdown with morpholino phosphorodiamidates and hydroxyprolylphosphono peptide nucleic acids. Nucleic Acids Res 33, 4914–4921 (2005).

41. E. Zamir, Z. Kam, A. Yarden, Transcription-dependent induction of G1 phase during the zebra fish midblastula transition. Mol Cell Biol 17, 529–536 (1997).

42. J. H. Hurley, ESCRTs are everywhere. EMBO J 34, 2398–2407 (2015).

43. S. H. Chan et al., Brd4 and P300 Confer Transcriptional Competency during Zygotic Genome Activation. Dev Cell 49, 867–881 e868 (2019).

44. N. Olivier et al., Cell lineage reconstruction of early zebrafish embryos using label-free nonlinear microscopy. Science 329, 967–971 (2010).

45. A. Caballe, J. Martin-Serrano, ESCRT machinery and cytokinesis: the road to daughter cell separation. Traffic 12, 1318–1326 (2011).

46. N. Chiaruttini et al., Relaxation of Loaded ESCRT-III Spiral Springs Drives Membrane Deformation. Cell 163, 866–879 (2015).

