## Supplementary Information for "Cytokinetic abscission is part of the mid-blastula transition switch in early zebrafish embryogenesis"

**This PDF file includes:**

- Supplementary methods
- Figures S1 to S5
- Table S1
- Legends for Movies S1 to S7
- SI References

**Other supplementary materials for this manuscript include the following:**

Movies S1 to S7

### Supplementary methods

#### Dextran injection

Embryos were injected with HiLyte 488-Tubulin as described in main methods. At 128 cell stage, embryos were dechorionized and mounted in an agar chamber with animal pole facing up. One cell was carefully injected with TAMRA Dextran 10,000 MW (20mg/ml in 0.2M KCL, D1816 Invitrogen) by iontophoretic injection. Injected embryo was placed at multi photon upright system and imaged over time (Olympus & Lavision-Biotec custom stand objective 20X, water, N.A 0.5). Time interval 1.5 minutes, 98 Z slices.

#### Western blot analysis

Injected and control embryos were collected at the indicated times and detached from their chorions using 0.5 mg/ml pronase (p-8811, Sigma Aldrich) in a dish coated with 1% agar. 50 embryos for each sample were collected and washed with ice-cold PBS. Embryos were then deyolked using a P200 pipet tip in ice-cold PBS and washed in ice-cold PBS. Embryos were homogenized in cholate extraction buffer (2% Sodium Deoxycholate, Sigma Aldrich D-6750, in 10mM Tris pH8.0), diluted in sample buffer and boiled (95°C for 5 minutes). Volume equivalent to 10 embryos were loaded in SDS-PAGE. Membranes were stained with the following antibodies: anti-VPS4 (1:1000, Cat # SAB4200025, Sigma-Aldrich), anti-CHMP4B (1:500, ab105767,abcam), anti GAPDH (1:1000, cat # G096, Applied Biological Materials) and rabbit or mouse-peroxidase secondary antibodies (1:10,000, catalog number 715-035-151 or 711-035-152, Jackson ImmunoResearch, West Grove, PA,). Bands intensities were quantified in ImageJ, using gel analysis plug-in.

#### Analysis of cell shape

To determine changes in cell shape, videos of embryos labeled with HiLyte 488-Tubulin during cycle 10 (MBT) were recorded for 8 time points at 2 minutes intervals. We manually selected cell triplets that included two connected cells and one non-connected cell, while avoiding triplets that included dividing cells in order to avoid quantifying cell shape that result from mitosis. Cell contours were manually annotated and their aspect ratios were calculated for all cells at all timepoints using SlideBook (version 6, Intelligent Imaging, Denver, CO). A cell's aspect ratio is the ratio between its major axis length and its minor axis length and is a measure the cells' elongated morphology. Aspect ratio of 1 indicates a rounded shape while larger values indicate more elongated shapes. We calculated the absolute difference in the aspect ratio between the cell pairs (connected and non-connected) in each triplet for each time point and averaged the absolute difference for each pair over time to reduce measurement noise (Fig S5*B*).

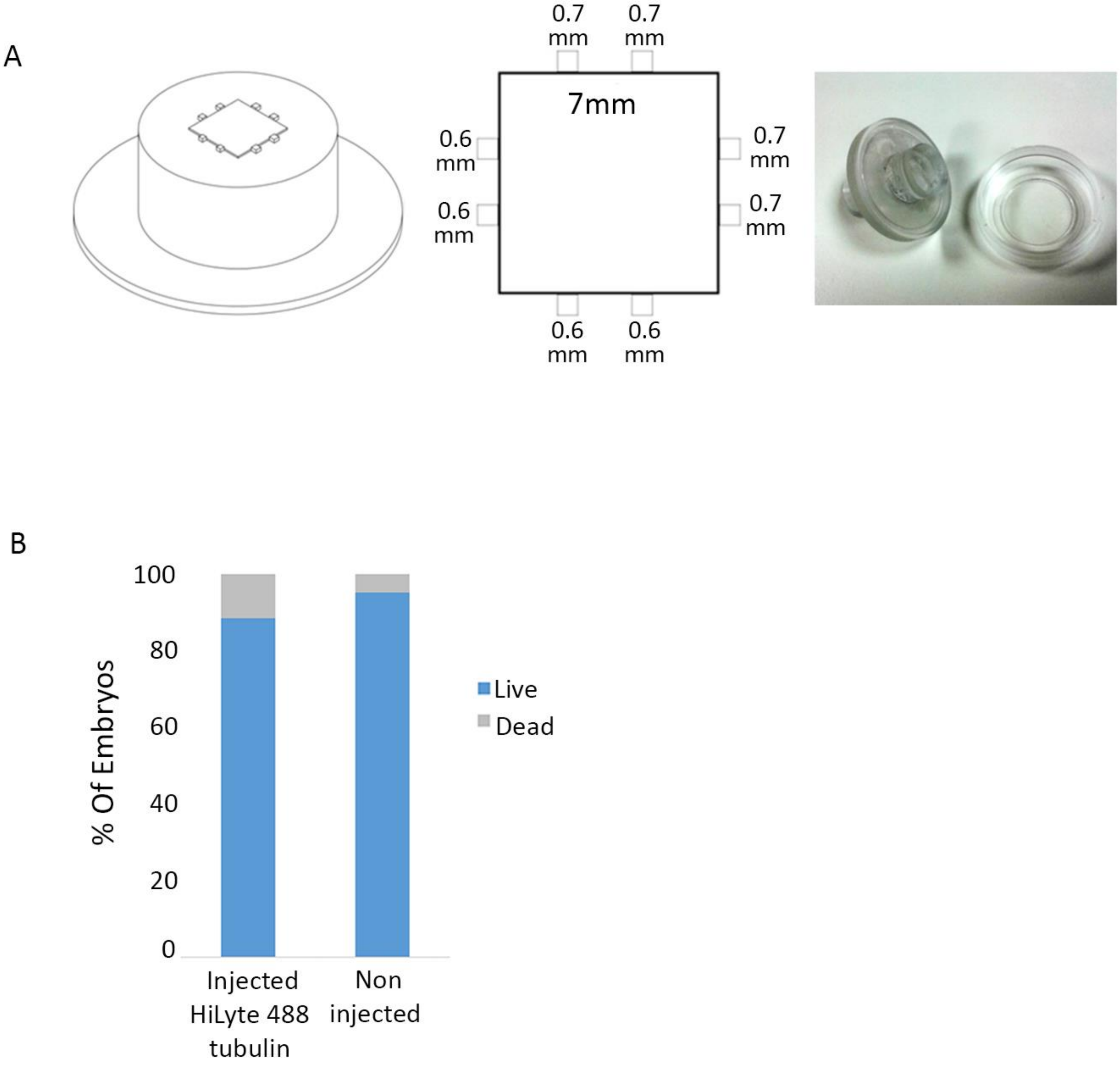


***Fig. S1: Experimental system***

(***A***) Survival of embryos injected or not injected at one-cell stage with HiLyte 488-Tubulin and phenol red was examined 24 hours post fertilization.  Injected embryos, 88.6% survival (n=60 embryos); Non injected embryos, 95.23% survival, n=30 embryos). Moreover, normal fish morphology was observed in injected embryos. These results indicate that the injection protocol of fluorescently labeled tubulin did not affect the overall developmental program of the fish. (***B***) Custom-made mold for preparation of agar chambers used for live imaging of zebrafish embryos. Mold was designed according to Wuhr et al (1) to make 8 wells in 0.6-0.7 $\mu m^2$ in 35mm glass bottom µ-dish. Chambers were prepared as described in materials and methods.

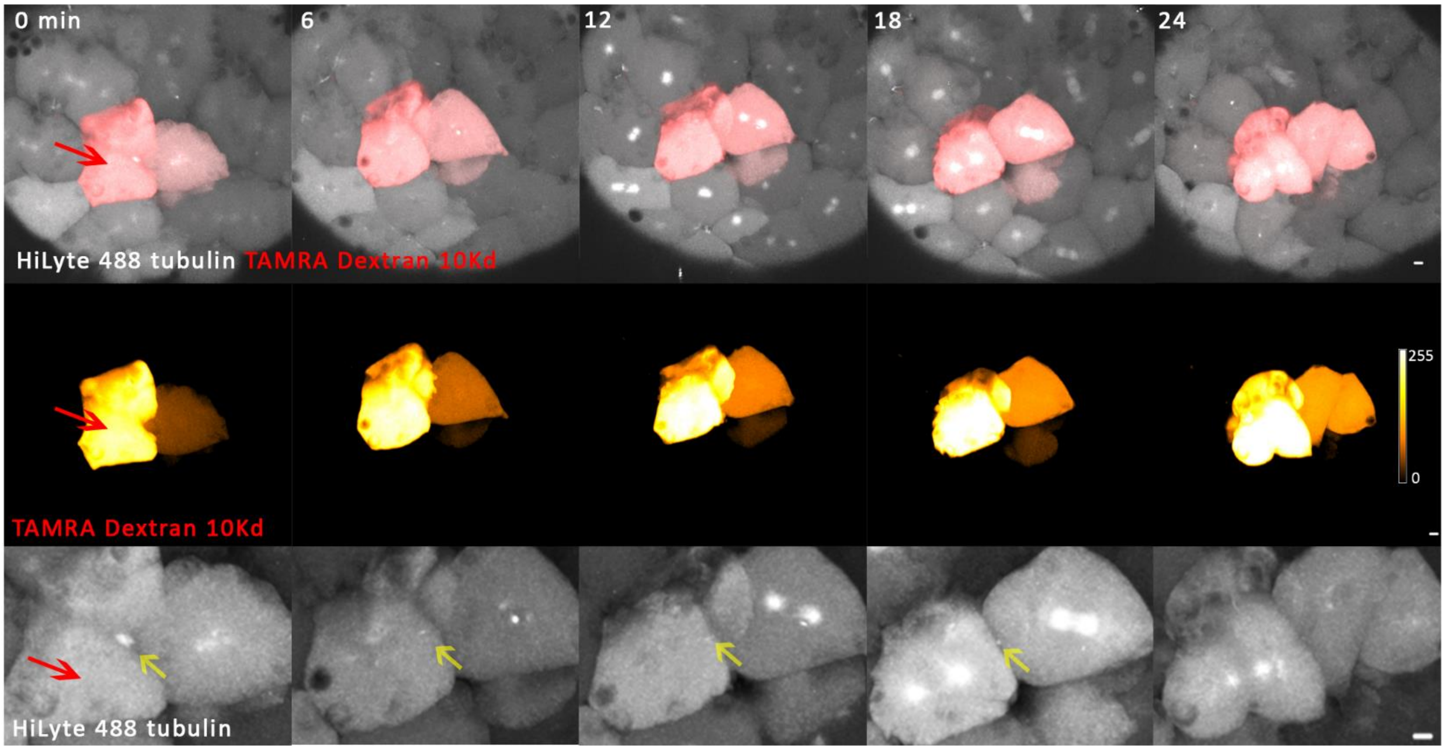


***Fig. S2: Dextran diffuses between clusters of cells interconnected by intercellular bridges***

128-cell embryo expressing HiLyte 488-Tubulin was injected with and TAMRA-Dextran, 10,000 MW as described in sup methods. Red arrow represents injection point. Cells were then recorded in 3D using a multi-photon microscope. Shown are maximum intensity projections images of 3D volumes (84 Z slices of 0.7µm) of selected time points, taken from the movie sequence (Movie S5). HiLyte 488-Ttubulin, white; TAMRA Dextran, red. Upper panel, merged images; middle panel, TAMRA-Dextran, color coded by intensity levels; bottom panel, HiLyte-488 Tubulin. Intercellular bridges are marked with yellow arrows. Note that dextran diffuses only between cells connected by intercellular bridges. Scale bar, 10µm

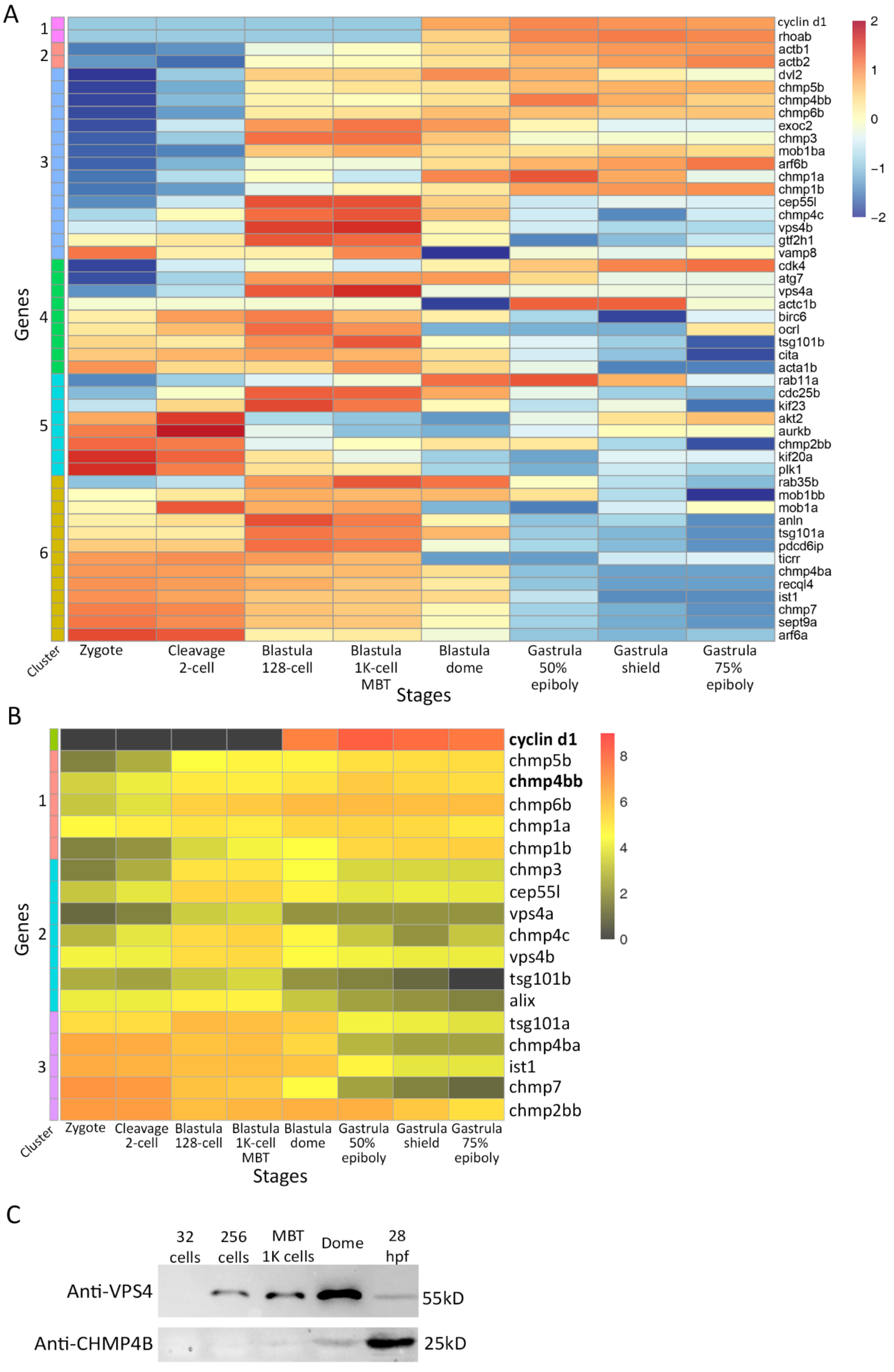

A
1
2
3
4
5
6
Genes
Cluster
cyclin d1
rhoab
actb1
actb2
dvl2
chmp5b
chmp4bb
chmp6b
exoc2
chmp3
mob1ba
arf6b
chmp1a
chmp1b
cep55l
chmp4c
vps4b
gtf2h1
vamp8
cdk4
atg7
vps4a
actc1b
birc6
ocrl
tsg101b
cita
acta1b
rab11a
cdc25b
kif23
akt2
aurkb
chmp2bb
kif20a
plk1
rab35b
mob1bb
mob1a
anln
tsg101a
pdcd6ip
ticrr
chmp4ba
recql4
ist1
chmp7
sept9a
arf6a
2
1
0
−1
−2
Zygote
Cleavage 2-cell
Blastula 128-cell
Blastula 1K-cell MBT
Blastula dome
Gastrula 50% epiboly
Gastrula shield
Gastrula 75% epiboly
Stages
B
1
2
3
Genes
Cluster
cyclin d1
chmp5b
chmp4bb
chmp6b
chmp1a
chmp1b
chmp3
cep55l
vps4a
chmp4c
vps4b
tsg101b
alix
tsg101a
chmp4ba
ist1
chmp7
chmp2bb
8
6
4
2
0
Zygote
Cleavage 2-cell
Blastula 128-cell
Blastula 1K-cell MBT
Blastula dome
Gastrula 50% epiboly
Gastrula shield
Gastrula 75% epiboly
Stages
C
32 cells
256 cells
MBT 1K cells
Dome
28 hpf
Anti-VPS4
55kD
Anti-CHMP4B
25kD

***Fig. S3: mRNA transcription levels of late Cytokinetic related genes at early development***

(***A***) RNA-seq profiles of 48 late cytokinetic related genes and *cyclin d1* across eight developmental stages of zebrafish embryos, downloaded from the data based established by White et al. are shown on a heatmap (2). Rows correspond to individual genes and columns correspond to developmental stages. Colors in heatmap indicate the expression levels of each gene across the samples, relative to its mean expression and trimmed to range [−2, 2]. Six clusters (Cluster 1-6, left bar color coding) were identified by hierarchical clustering, clusters were sorted manually, and genes within each cluster were sorted by their presented (relative) expression in the Zygote stage. (***B***) RNA- seq profiles of ESCRT genes (raw data). RNA sequencing data of zebrafish embryos from eight developmental stages is used to present the expression levels of *cyclin d1* and 17 ESCRT genes. Log2-transformed FPKM levels of the genes are presented on a heatmap, genes are sorted according to hierarchical clustering as presented in Fig 3B. Rows correspond to individual genes and columns correspond to developmental stages. Colors represent the expression levels of each gene across the samples [log2(FPKM)]. Expression values are trimmed to range [0, 9]. (***C***) Expression levels of ESCRT proteins at different embryonic stages as determined by western blot analysis. Wild-type embryos were lysed at indicated stages and subjected to western blot analysis using anti-VPS4 and anti-CHMP4B antibodies. Volumes equivalent to 10 embryos were loaded in each lane. The results are in agreement with the data presented in heatmap for mRNA expression. Vps4 is not expressed at 32-cell stage, and its levels gradually increase until dome stage while Chmp4b is not expression prior to MBT (n=3).

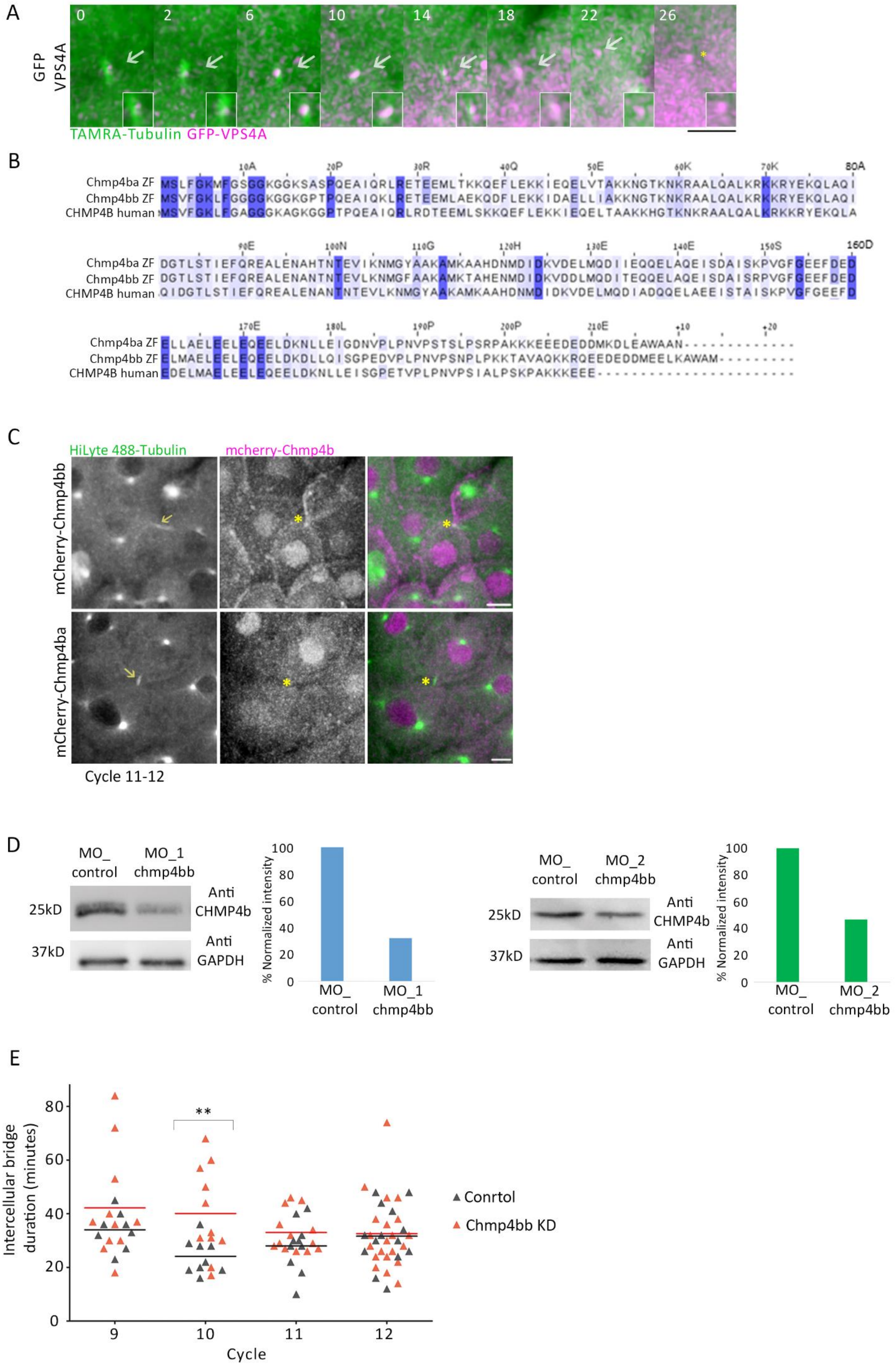

A
GFP VPS4A
0
2
6
10
14
18
22
26
TAMRA-Tubulin GFP-VPS4A
B
10A
20P
30R
40Q
50E
60K
70K
80A
Chmp4ba ZF MSLFGKMFGSGGKGGKSASPQEAIQRLRETEEMLTKKQEFLEKKIEQELVTAKKNGTKNKRAALQALKRKKRYEKQLAQI
Chmp4bb ZF MSVFGKLFGGGGKGGKGPTPQEAIQKLRETEEMLAEKQDFLEKKIDAELLIAKKNGTKNKRAALQALKRKKRYEKQLAQI
CHMP4B human MSVFGKLFGAGGGKAGKGGPTPQEAIQRLRDTEEMLSKKQEFLEKKIEQELTAAKKHGTKNKRAALQALKRKKRYEKQLA
90E
100N
110G
120H
130E
140E
150S
160D
Chmp4ba ZF DGTLSTIEFQREALENAHTNTEVIKNMGYAAKAMKAAHDNMDIDKVDELMQDIIEQQELAQEISDAISKPVGFGEEFDED
Chmp4bb ZF DGTLSTIEFQREALENANTNTEVLKNMGFAAKAMKTAHENMDIDKVDDLMQDITEQQELAQEISDAISRPVGFGEEFDED
CHMP4B human QIDGTLSTIEFQREALENANTNTEVLKNMGYAAKAMKAAHDNMDIDKVDELMQDIADQQELAEEISTAISKPVGFGEEFD
170E
180L
190P
200P
210E
+10
+20
Chmp4ba ZF ELLAELEELEQEELDKNLLEIGDNVPLPNVPSTSLPSRPAKKKEEEDEDDMKDLEAWAAN
Chmp4bb ZF ELMAELEELEQEELDKDLLQISGPEDVPLPNVPSNPLPKKTAVAQKKRQEEDEDDMEELKAWAM
CHMP4B human EDELMAELEELEQEELDKNLLEISGPETVPLPNVPSIALPSKPAKKKEEE
C
HiLyte 488-Tubulin
mcherry-Chmp4b
mCherry-Chmp4bb
mCherry-Chmp4ba
Cycle 11-12
D
MO_ control
MO_1 chmp4bb
25kD
37kD
Anti CHMP4b
Anti GAPDH
% Normalized intensity
100
80
60
40
20
0
MO_ control
MO_1 chmp4bb
MO_ control
MO_2 chmp4bb
25kD
37kD
Anti CHMP4b
Anti GAPDH
% Normalized intensity
100
80
60
40
20
0
MO_ control
MO_2 chmp4bb
E
Intercellular bridge duration (minutes)
80
60
40
20
0
**
9
10
11
12
Cycle
Conrtol
Chmp4bb KD

***Figure S4: ESCRT proteins in zebrafish embryogenesis***

(***A***) Live imaging of VPS4 arrival to the intercellular bridge of cells in embryos at blastula stage. Embryos were injected with mRNA that encodes for VPS4A component fused to GFP (magenta) and with HiLyte 488-Tubulin (green). Shown are maximum intensity projection images (10 Z slices of 0.7 µm intervals) taken from a representative cell imaged in embryos at blastula stage. VPS4A localized to the intercellular bridge (gray arrow) and was detected at the midbody remnant post abscission (yellow asterisk). Time 0 represents the time of bridge formation. Zoomed-in images of the midbody are shown as insets in each frame. data was obtained from 2 embryos and a total of 10 intercellular bridges. Scale bar 10 µm (***B***) Multiple sequence alignment of human CHMP4B (NP_789782.1), zebrafish Chmp4ba (NP_956489.1) and zebrafish Chmp4bb (NP_998622.1). (***C***) Visualizing the two CHMP4B homologs in cytokinetic cells of zebrafish embryos. Embryos were injected with mRNA that encodes to Chmp4bb (upper panel) and Chmp4ba (bottom panel) fused to mCherry (magenta) and with HiLyte 488-tubulin (green). Shown are maximum intensity projection images (10 Z slices of 0.9-1 µm intervals) taken from a representative cell imaged in embryos at cycle12$^{th}$. Intercellular bridges are indicated by arrows.  Chmp4bb localized to the intercellular bridge while Chmp4ba exhibited diffused pattern and could not be detected at the intercellular bridge (yellow arrow). Scale bar 10µm. (***D***) Depleting Chmp4bb levels using morpholino sequences. Embryos were injected with MO_control, MO_1chmp4bb or MO_2chmp4bb, lysed 24 hours post fertilization and subjected to western blot analysis using CHMP4B and GAPDH antibodies. Volume equivalent to 10 embryos was loaded for each treatment. Chmp4bb levels were reduced by 67.9% in MO_1chmp4bb and by 53.4% in MO_2chmp4bb compared to MO_control embryos. MO_1chmp4bb was therefore used for Chmp4bb depletion experiments. (***E***) Duration of intercellular bridges in Chmp4bb depleted embryos. Zebrafish embryos were injected with MO_1chmp4bb oligo and HiLyte 488-Tubulin and intercellular bridges were imaged in embryos at cycle 9 to 12. Intercellular bridge duration was measured as described in material and methods. Averaged bridge duration in Chmp4bb depleted embryos: cycle 9, 42.18± 19.9 minutes (n=11); cycle 10, 40.09±16.82 minutes (n=11); cycle 11 x=33± 7.48 (n=13); cycle 12, 32.57± 13.34 minutes (n=21). Averaged bridge duration in control embryos: cycle 9, 34±6.96 minutes (n=8); cycle 10, 24.11±6.47 minutes (n=9); cycle 11, 28± 9.56 (n=10); cycle 12, 31.64±10.88 minutes (n=14). ± indicate S.D. n indicates number of bridges. Data was obtained from 8 embryos.

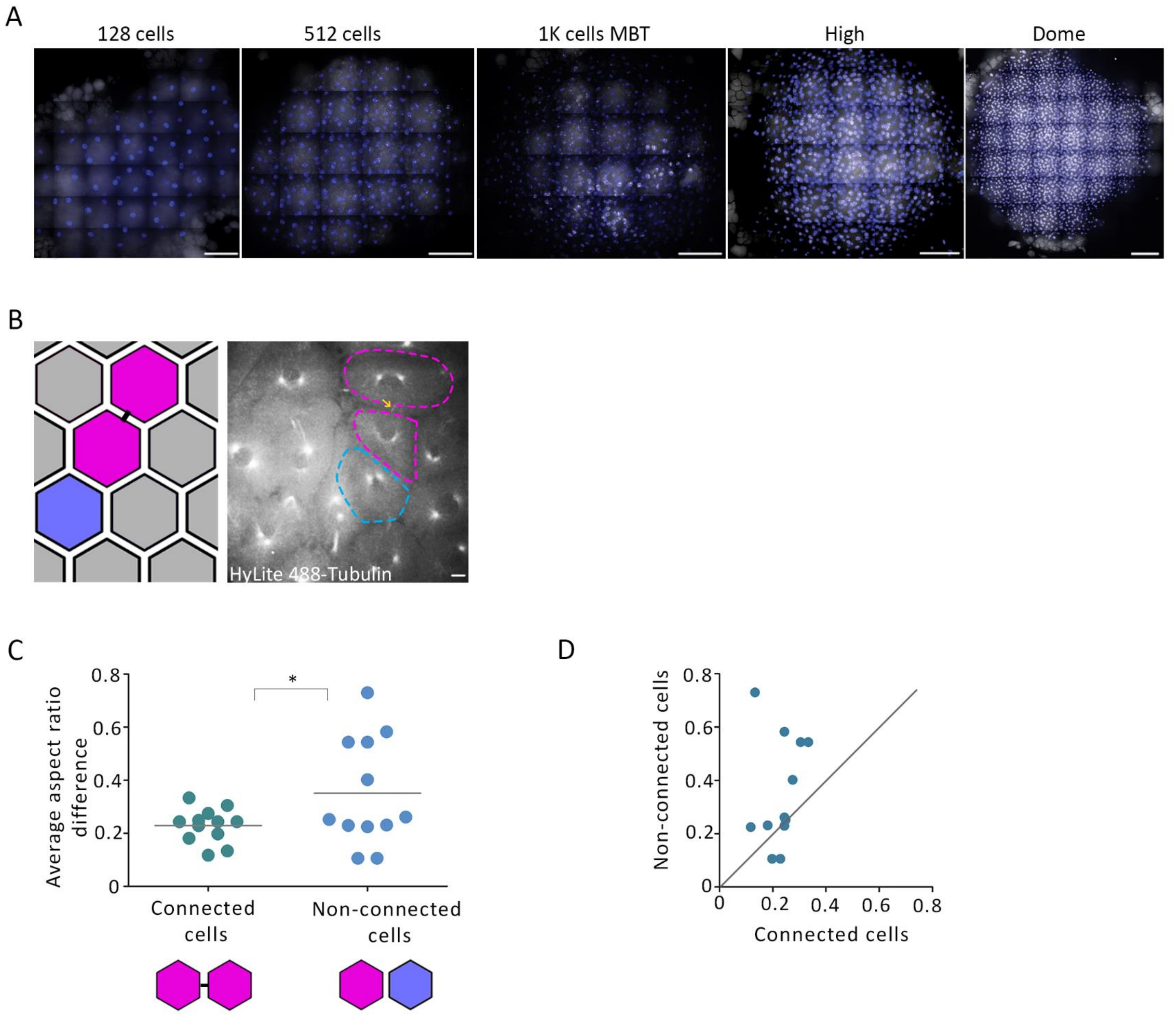


***Figure S5: Similarity between connected cells in the embryo***

(***A***) Characterizing transcription in zebrafish embryos at blastula stage. Embryos were treated and imaged as described in Fig 4. Shown are nuclear and 5-EU labeling of the entire embryos presented in Fig 4B. Scale bar 100 µm. (***B-D***) quantitative analysis of the similarity in cell shape observed in pairs of cells connected with intercellular bridges and in pairs of non-connected neighboring cells (determined based on Tubulin labeling as described in supplementary methods). (***B***) Representation of the analysis. Cell triplets included one cell pair connected with a bridge (magenta) and another pair with no bridge connection (blue). Left, schematic representation; right, a representative cell triplet. Scale bar 10μm. (Movie S7) (***C***) The absolute difference in the mean aspect ratio between connected cell pairs (magenta- magenta) non-connected cell pairs (magenta-blue) (*supplementary methods*). The absolute difference in aspect ratios between connected cells 0.23±0.05 was significantly lower than in non-connected neighboring cells (0.35±0.19, *t*-test, $t_{0.05,16}$=1.915, *p*=0.0368). Data was obtained from 5 embryos and total of 12 cell triplets of cells, over 8 time points. (***D***) The matched absolute difference in the aspect ratio between connected cell pairs and their corresponding non-connected pair. The diagonal Y = X is a guide to the eye, match observations above the line indicate larger aspect ratio deviation in non-connected pairs versus their matched connected pair. Wilcoxon matched-pairs signed rank rejects the null hypothesis that the difference between the matched connected- and non-connected pairs is distributed around zero (n=12, Z=-50, *p*=0.026). ± indicate S.D

*Table S1. Primers and oligos*

| Gene | Method | Primer | Restriction enzyme |
|---|---|---|---|
| cep55l zf Fw | restriction enzyme cloning to mCherry C1 vector | 5'-TGTGGAGATCTATGGCGGCGAAGGG-3' | BglII |
| cep55l zf Rv | | 5'-TGGCGGGTACCTTAGGTGAAGCAGTAGTCG-3' | KpnI |
| tsg101 zf FW | | 5'-TGTGGCTCGAGCTATGGCTGTTGTCAACGAAG-3' | XhoI |
| tsg101 zf RV | | 5'-TGGCGGGTACCTCAGTATAGATCACTAAGTCCAGC-3' | KpnI |
| chmp4bb zf Fw | | 5'-TGTGGCTCGAGCTATGTCTGTATTCGGCAAATTGTTTGGC-3' | XhoI |
| chmp4bb zf Rv | | 5'-TGGCGGGATCCTCACATGGCCCATGCTTTGAGTTCC-3' | BamHI |
| chmp4ba zf Fw | | 5'-GCTGCTCTCGAG CTATGTCTTTGTTCGGGAAG-3' | XhoI |
| chmp4ba zf Rv | | 5'-GCTGCTGGATCCTTAATTGGCAGCCC-3' | BamHI |
| mcherry Fw | Gibson assembly into pcs2+ vector | 5'-GCTACTTGTTCTTTTTGCAGATGGCCTCCTCCGAGGAC-3' | |
| mcherry-cep55 Rv | | 5'-TAGAGGCTCGAGAGGCCTTGTTAGGTGAAGCAGTAGTCGAGG-3' | |
| TSG101 zf OL pCS2 GA RV | | 5'-TAGAGGCTCGAGAGGCCTTGTCAGTATAGATCACTAAGTCCAGCAGTC-3' | |
| mcherry-chmp4bb Rv | | 5'-TAGAGGCTCGAGAGGCCTTGTCACATGGCCCATGCTTT-3' | |
| mcherry-chmp4ba Rv | | 5'-CACCGGCGCCTCCGGACTCAGATCTATGTCTTTGTTCGGGAAGATG-3' | |
| mcherry-chmp4bb Rv | | 5'-AGTTCTAGAGGCTCGAGAGGCCTTGTTAATTGGCAGCCCAAGC-3' | |
| MO_1chmp4bb | morpholino oligos for microinjection | 5'-ATTTGCCGAATACAGACATATTGTT-3' | |
| MO_2chmp4bb | | 5'-AATCCCTGTTTGTCTTCTAGACACC-3' | |
| MO_standard control | | 5'-CCTCTTACCTCAGTTACAATTTATA-3' | |

**Movie S1. Visualizing mitosis in live zebrafish embryos**

Embryo was injected with HiLyte-488 tubulin and imaged live during early development (cycles 5-15) using a 20X N.A 0.8 objective to visualize mitosis in the embryo. Maximum intensity projection (30 Z slices at 1 µm) are shown of 4minutes intervals. Scale bar 10 µm

**Movie S2. Visualizing intercellular bridges in live zebrafish embryos**

Embryo was injected with HiLyte-488 tubulin and imaged during early development (cycles 9-13) using a 40X N.A 1.3 objective to visualize intercellular bridges at the embryo. Maximum intensity projection (30 Z slices at 1 µm) are shown. Scale bar 10 µm

**Movie S3. Intercellular bridge duration**

Embryo was injected with HiLyte 488 tubulin (green) and mRNA encoding to mCherry-CEP55 (red) and imaged live using a 40X objective to track the duration of intercellular bridge in the embryo. Maximum intensity projection (29 Z slices at 0.8 µm) are shown. Scale bar 5 µm

**Movie S4. Persistence on intercellular bridges in early embryos**

Embryo was injected with HiLyte 488 tubulin (green) and mRNA encoding to mCherry-CEP55 (red). 1$^{st}$ bridge (white arrow) was formed in the end of cycle 8. At the end of cycle 9, a 2$^{nd}$ bridge (yellow arrow) is formed while the1$^{st}$ bridge was yet to be resolved. Scale bar 10 µm

**Movie S5. Dextran diffusion between clusters of cells inter-connected by intercellular bridges**

TAMRA-Dextran 10,000 MW (red) was injected to a single cell in a 128 cells embryo that was pre injected with HiLyte-488 tubulin (white). Dextran diffused from the injected cell to cells connected by intercellular bridges but was excluded from neighboring cells. Scale bar 10 µm

**Movie S6. Bridge duration Control and knock down embryos**

(A) Embryo was injected with HiLyte 488 tubulin (white) and MO_control and imaged live using a 40X objective to track the duration of intercellular bridge (yellow arrow) in the embryo. Maximum intensity projection (12 Z slices at 1 µm) are shown. Scale bar 5 µm

(B) Embryo was injected with HiLyte 488 tubulin (white) and MO_1chmp4bb and imaged live using a 40X objective to track the duration of intercellular bridge (yellow arrow) in the embryo. Maximum intensity projection (20 Z slices at 1 µm) are shown. Scale bar 5 µm

**Movie S7. Changes in cells shape**

Tracking cell shape changes in embryos at MBT. White, HiLyte-488 tubulin; Blue, cell masks used for aspect ratio calculations. Scale bar 10 µm

REFERENCES


1. M. Wuhr, N. D. Obholzer, S. G. Megason, H. W. Detrich, 3rd, T. J. Mitchison, Live imaging of the cytoskeleton in early cleavage-stage zebrafish embryos. *Methods Cell Biol* **101**, 1-18 (2011).
2. R. J. White *et al.*, A high-resolution mRNA expression time course of embryonic development in zebrafish. *Elife* **6** (2017).
